# Leveraging autophagy and pyrimidine metabolism to target pancreatic cancer

**DOI:** 10.1101/2025.05.29.656904

**Authors:** Suzanne Dufresne, Ramya S Kuna, Kristiana Wong, Anvita Komarla, Louis R Parham, Celina Shen, Payel Mondal, Adelaida Estrada-Cardenas, Angelica Rock, Joel Rosada-Encarnación, Amanda Cyril, Ginevra Doglioni, Dhanya R. Panickar, Nicole A. Bakas, Douglas J. Sheffler, Allison Limpert, Fabiana Izidro Layng, Kristina L Peck, Alexandra Fowler, Sonja N. Brun, Tammy J. Rymoff, Andrew M Lowy, Dannielle Engle, Herve Tiriac, Reuben J. Shaw, Nicholas D.P. Cosford, Christian Metallo, Christina G. Towers

## Abstract

Autophagy inhibitors are promising compounds to treat pancreatic ductal adenocarcinoma (PDA) but their efficacy in patients is unclear, highlighting a need to understand mechanisms of resistance. We used a novel approach to uncover metabolic adaptations that bypass autophagy inhibition. Utilizing PDA cells with acquired resistance to different autophagy inhibitors, we found that severe autophagy depletion induces metabolic rewiring to sustain TCA intermediates and nucleotides for biosynthesis. Long-term autophagy inhibition results in altered pyruvate metabolism likely regulated by lower pyrimidine pools. Cells adapting to loss of autophagy preferentially salvage pyrimidines to replenish these pools instead of synthesizing them de novo. Exploiting this metabolic vulnerability, we found that acquired resistance to autophagy inhibition promotes increased salvage and therefore sensitivity to pyrimidine analogues, including gemcitabine and trifluridine/tipiracil leading to combinatory effects with autophagy inhibitors and pyrimidine analogs. These studies provide mechanistic insight defining how autophagy inhibition can be leveraged to treat pancreatic cancer.

## INTRODUCTION

Pancreatic ductal adenocarcinoma (PDA) is a highly aggressive form of pancreatic cancer with limited treatment options. This devastating disease undergoes heavy metabolic rewiring during the course of tumor progression and in response to treatment. Therefore, many efforts are tring to leverage metabolic vulnerabilities to develop better therapeutic strategies, although thus far with little clinical success.

Over the past decade, the cellular recycling process, autophagy, has emerged as a promising therapeutic target for this disease ^1–4^. Autophagy is a catabolic process that degrades and recycles damaged cellular material by trafficking cargoes to the lysosome ^5–7^. It is tightly regulated and induced by extracellular and intracellular stress stimuli such as nutrient deprivation and hypoxia. As a coordinated stress response, autophagy maintains metabolic homeostasis, protein turnover, and organelle quality control – thereby supporting cancer cell survival. Oncogene activation, including *KRAS* mutations – present in more than 90% of PDA, increases basal autophagy levels^8–10^. Aberrantly high levels of autophagy provide a survival advantage to PDA cells in nutrient-deprived and hypoxic conditions characteristic of pancreatic cancer^11^. Autophagy-mediated recycling provides a variety of substrates to fuel nearly all aspects of central carbon metabolism^12,13,10,14–18^. In addition, autophagy recycles nucleic acids into nucleotide building blocks to preserve nucleotide homeostasis^19,20^, a necessary metabolic trait for the maintenance of high proliferative rates of PDA cells^21^.

Numerous studies have shown that perturbation of autophagy decreases PDA cell survival across in vitro and in vivo models. For example, deletion of the core autophagy gene Atg5 or expression of a dominant negative mutant of *Atg4b* in the classic *Kras^LSL-G12D/+^*, *Tp53^flox/+^*, *Pdx1-Cre* (KPC) PDA model results in slower tumor growth and increased survival^22,23^. Therefore, autophagy represents an attractive potential target for this devastating disease. Chloroquine (CQ) and hydroxychloroquine (HCQ) are FDA-approved drugs, initially used to treat malaria, that block autophagy and lysosomal function. Although chloroquine derivatives are potent autophagy inhibitors in cultured cells, low in vivo bioavailability along with low specificity limit their therapeutic potential. Therefore, recent efforts have led to the development of more specific autophagy inhibitors including compounds that target the upstream Unc-51 Like Autophagy Activating Kinases 1 and 2 (ULK1/2) as well as other autophagy-regulating enzymes ^24–26^.

In cancer patients, over 60 clinical trials combining CQ or HCQ with standard of care therapies have launched^27^. Despite promising preclinical findings, the results from clinical trials have been disappointing, likely caused by a compilation of inherent and acquired therapeutic resistance along with inadequate drug bioavailability^28^. This is supported by previous findings from our lab showing that lung and breast cancer cells that are initially dependent on autophagy and sensitive to genetic deletion of core autophagy genes like *ATG7* or *RB1CC1* (*FIP200*) can adapt over time and become autophagy-independent^29,30^. However, whether PDA cells can acquire resistance to pharmacological blockade of autophagy inhibition remains unknown. Prior studies have explored different drug combinations that could improve autophagy inhibition to treat PDA including MAP Kinase pathway inhibition^31,32,25,26^, targeting insulin like growth factor 1 receptor (IGF1R)^33^, and autophagy related processes like macropinocytosis^34^, each demonstrating increased efficacy over single agent use of autophagy inhibitors. Nonetheless, these efforts have not yet resulted in better, clinically relevant, therapeutic strategies to target autophagy in PDA.

In this study, we took a unique approach to investigate the metabolic alterations and adaptations PDA cells use to acquire resistance to autophagy inhibitors. For the first time, we generated PDA cells resistant to either the lysosomal inhibitor, HCQ, or the ULK1/2 inhibitor, MRT68921. Cells with acquired resistance maintain autophagy inhibition but undergo metabolic rewiring to adapt and survive. The resistant cells have altered pyruvate metabolism and pyrimidine metabolism, resulting in increased pyrimidine salvage. We exploited this vulnerability and found that acquired resistance to autophagy inhibitors resulted in increased salvage of pyrimidine analogues and therefore a dramatic sensitivity to standard-of-care drugs like gemcitabine as well other clinically relevant pyrimidine analogues like the thymidine analogue, trifluridine. Together, our findings provide mechanistic insight into metabolic alterations that occur in reponse to autophagy inhibition and indicate that combining autophagy inhibitors and pyrimidine analogs represents a promising therapeutic strategy for PDA.

## RESULTS

### PDA cells that acquire resistance to autophagy inhibition have blocked autophagy

To interrogate if and how PDA cells acquire resistance to pharmacological autophagy inhibition, we used murine PDA cell lines derived from the well-established Kras^LSL-G12D/+^, p53^flox/+^, Pdx1-Cre (KPC) mouse model including FC1199 (male) and FC1245 (female) cells. The cells were exposed to increasing doses of the autophagy inhibitor hydroxychloroquine (HCQ) or the ULK1/2 kinase inhibitor MRT68921 (MRT), starting at the IC50 dose for each drug (Figure 1A). The cells were continuously passaged in the drug dose until they resumed active proliferation and minimal cell death, at which point a new IC50 was determined. The cells were then treated with the new IC50 dose, and this cycle was repeated 4-6 times until a 5-10-fold increase in the IC50 was reached.

**Figure 1.**
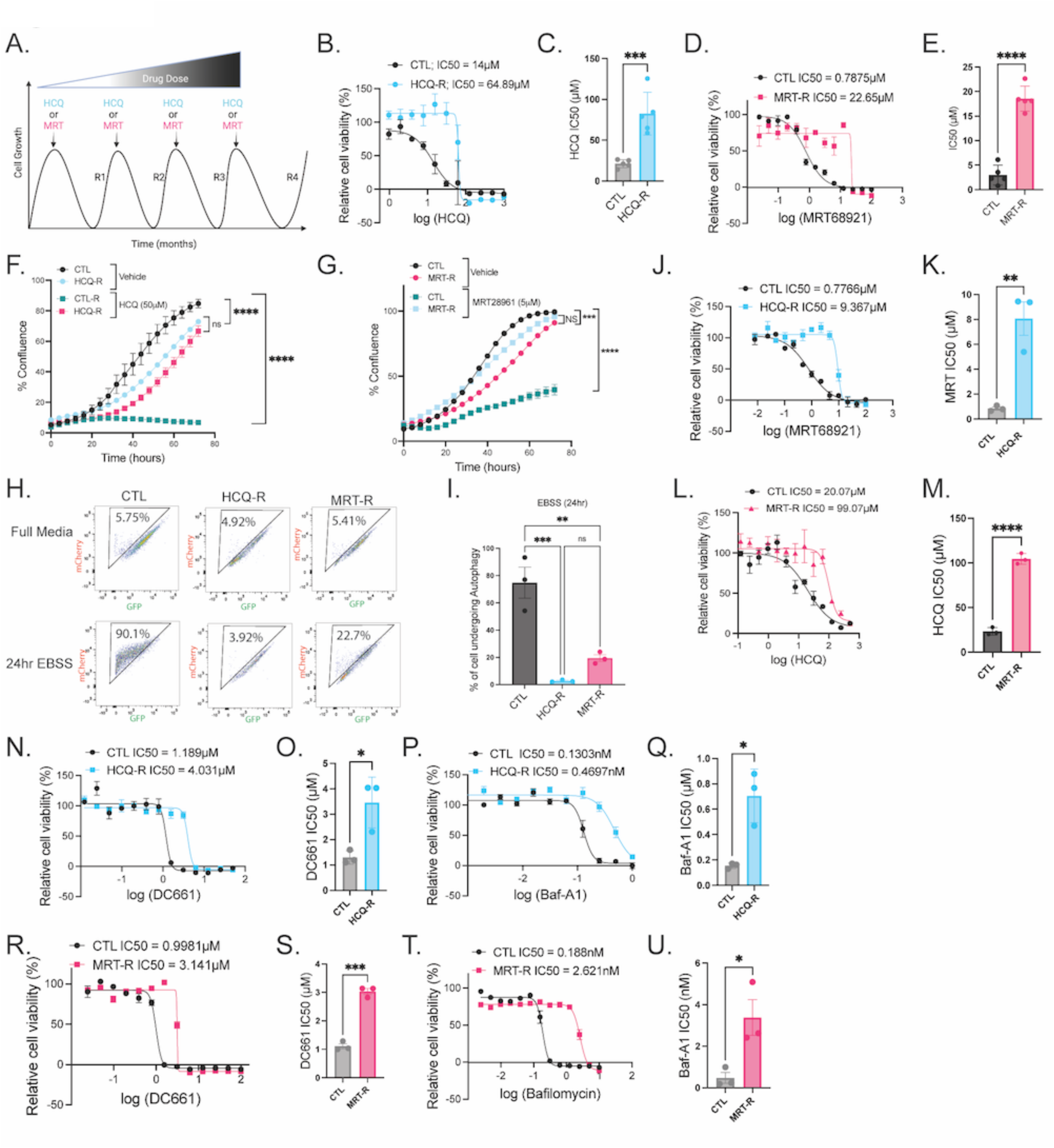
PDA cells that acquire resistance to autophagy inhibition have blocked autophagy. (A) Schematic overview of the process used to make resistant cells. (B-C) MTT cell viability assay in FC1199 CTL and HCQ-R cells treated with HCQ for 72h (B) and calculated IC50 values (C). (D-E) MTT cell viability assay in FC1199 CTL and MRT-R cells treated with MRT68921 for 72h (D) and calculated IC50 values (E). (F-G) Incucyte live cell imaging in FC1199 CTL, HCQ-R or MRT-R cells treated with HCQ or MRT68921 as indicated for 3 days. (H-I) FC1199 CTL, HCQ-R and MRT-R cells transfected with the mCherry-EGFP-LC3 tandem, starved for 24hrs in EBSS, or treated with Baf A1 (20nM) for 24hrs. mCherry/eGFP fluorescence was measured by flow cytometry (H) and autophagy induced under EBSS starvation was determined (I). (J-K) MTT cell viability assay in FC1199 CTL and HCQ-R cells treated with MRT68921 for 72h (J) and calculated IC50 values (K). (L-M) MTT cell viability assay in FC1199 CTL and MRT-R cells treated with HCQ for 72h (L) and calculated IC50 values (M). (N-Q) MTT cell viability assay in FC1199 CTL and HCQ-R cells treated with DC661 (N-O) or Baf A1 (P-Q) for 72h to determine the IC50 values. (R-U) MTT cell viability assay in FC1199 CTL and MRT-R cells treated with DC661 (N-O) or Baf A1 (P-Q) for 72h to determine the IC50 values. Data are represented as a mean ± SEM. Data on panels B, D, F-G, J, L, N, P, R and T show n=3 technical replicates representative of n=3 biological replicates. Data on panels C, E, I, K, M, O, Q, S and U represent n=3 biological replicates. ^∗^ indicates p<0.05, ^∗∗^ p<0.001, ^∗∗∗^ p <0.0001, ^∗∗∗∗^ p<0.00001 determined by two-tailed Student’ *t* test (C, E, K, M, O, Q, S and U), two-way repeated measures ANOVA with Tukey’s post hoc test (F-G) and one-way ANOVA with Tukey’s post hoc test (I). MRT68921 abbreviated as MRT.

Cell viability assays were used to confirm the level of resistance, and cells exposed to HCQ had a 5-fold increase in IC50 dose, hereafter referred to as HCQ-R cells, compared to CTL cells passaged without drug alongside the resistant cells (Figure 1B-C, S1A-B). Similarly, cells exposed to MRT68921, showed a 6-fold increase in IC50 dose, hereafter referred to as MRT-R (Figure 1D-E). Drug resistance was confirmed by cell proliferation assays using Incucyte live-cell imaging as well as colony formation assays. CTL cells showed an abrupt decrease in proliferation when exposed to HCQ or MRT, while the HCQ-R and MRT-R cells showed no change in viability (Figure 1F-G, S1C-D). The resistant cell lines actively proliferate while in drug, but their basal growth rates are slightly slower than their matched CTL counterparts. To assess whether this resistance can be obtained in other PDA cell lines, we exposed the human PDA cell line Panc-1 to increasing doses of HCQ and obtained similar resistance (Figure S1E).

To determine if autophagy is blocked in the HCQ-R and MRT-R cells, autophagy flux was assessed with the mCherry-GFP-LC3 tandem fluorescent reporter^35,36^. LC3-II decorates autophagosomes and is degraded when autophagosomes fuse with lysosomes, which possesses an acidic lumen. GFP is pH sensitive, and the fluorescence is quenched in the lysosome, whereas mCherry is pH stable. Ratiometric flow cytometry can be used to quantify the ratio of mCherry over GFP as a direct measure of autophagic flux. After expressing mCherry-GFP-LC3 in CTL, HCQ-R and MRT-R cells, we found the CTL cells responded to 24hr amino acid starvation in Earl’s Balanced Salt Solution (EBSS) with a robust increase in autophagic flux. However, the HCQ-R and MRT-R cells were unable to induce autophagy in these conditions (Figure 1H-I) highlighting that adaptation to the autophagy inhibitors does not reinstate autophagic flux.

Next, we tested how the resistant cells respond to other autophagy inhibitors. Cell viability and clonogenic assays revealed that the HCQ-R cells are also resistant to MRT68921 while the MRT-R cells are resistant to HCQ (Figure 1J-M, S1D, S1F-G). Moreover, both the HCQ-R and MRT-R cells are resistant to other autophagy inhibitors including the chloroquine derivative, DC661, and the lysosome inhibitor, Bafilomycin-A1 (Figure 1N-U). While both HCQ and MRT68921 have some off-target effects, these results show that the acquired resistance is on-target and autophagy is impaired in the HCQ-R and MRT-R cells. Importantly, the resistant cells proliferate indicating that they have adapted to bypass autophagy.

### PDA cells resistant to autophagy inhibition rewire their metabolism and decrease pyruvate decarboxylation

To assess the modifications necessary to bypass pharmacological blockade of autophagy, we performed RNA sequencing on the autophagy inhibition resistant FC1199 cells and drug naïve CTL cells. We compared differentially expressed genes (DEGs) and the corresponding Gene Ontology (GO) pathways from the two resistant cell line lines along with autophagy deficient *Atg7* KO cells (as a positive control for autophagy-inhibited pathways). There was a significant overlap in altered pathways suggesting that some mechanisms of resistance are shared between these two cell lines (Figure 2A, S2A-B). Within these shared GO terms, numerous metabolism-related pathways were altered, both positively and negatively, compared to CTL cells (Figure 2B). To better discern the metabolic changes in the resistant cells, we performed metabolomic assays (Figure S2C). HCQ-R and MRT-R cells showed significantly higher levels of pyruvate, but decreased citrate levels compared to drug naïve CTL cells (Figure S2C). Moreover, the HCQ-R cells had increased aspartate (Figure S2C). To understand the fate of pyruvate in the resistant cells, we performed a [U-^13^C_6_] glucose trace using gas chromatography-mass spectrometry (GC-MS) and assessed labeling in TCA intermediates. We confirmed increased pyruvate and decreased citrate abundances in the resistant cells (Figure 2C-D, S2D-I). There was also a consistent decrease in ⍺-ketoglutarate (⍺KG) in the HCQ-R and MRT-R lines (Figure S2J-K). These results could be due to altered pyruvate dehydrogenase complex (PDH) function. Once imported into mitochondria, glycolytic-derived pyruvate can undergo pyruvate decarboxylation when processed by PDH into acetyl-CoA. Acetyl-CoA then condensates with oxaloacetate (OAA) to generate citrate which enters the tricarboxylic acid (TCA) cycle. In the presence of [U-^13^C_6_] glucose, M3 pyruvate undergoes pyruvate decarboxylation leading to M2 citrate (Figure 2E). We found a decrease in the M2 Citrate over M3 Pyruvate ratio in the autophagy inhibition resistant cells (Figure 2F), indicative of decreased pyruvate decarboxylation. Datamining gene expression across human PDA cell lines through Depmap also revealed a positive correlation between autophagy-lysosome gene signatures^37^ and regulation of PDH which is responsible for pyruvate decarboxylation (Figure 2G).

**Figure 2.**
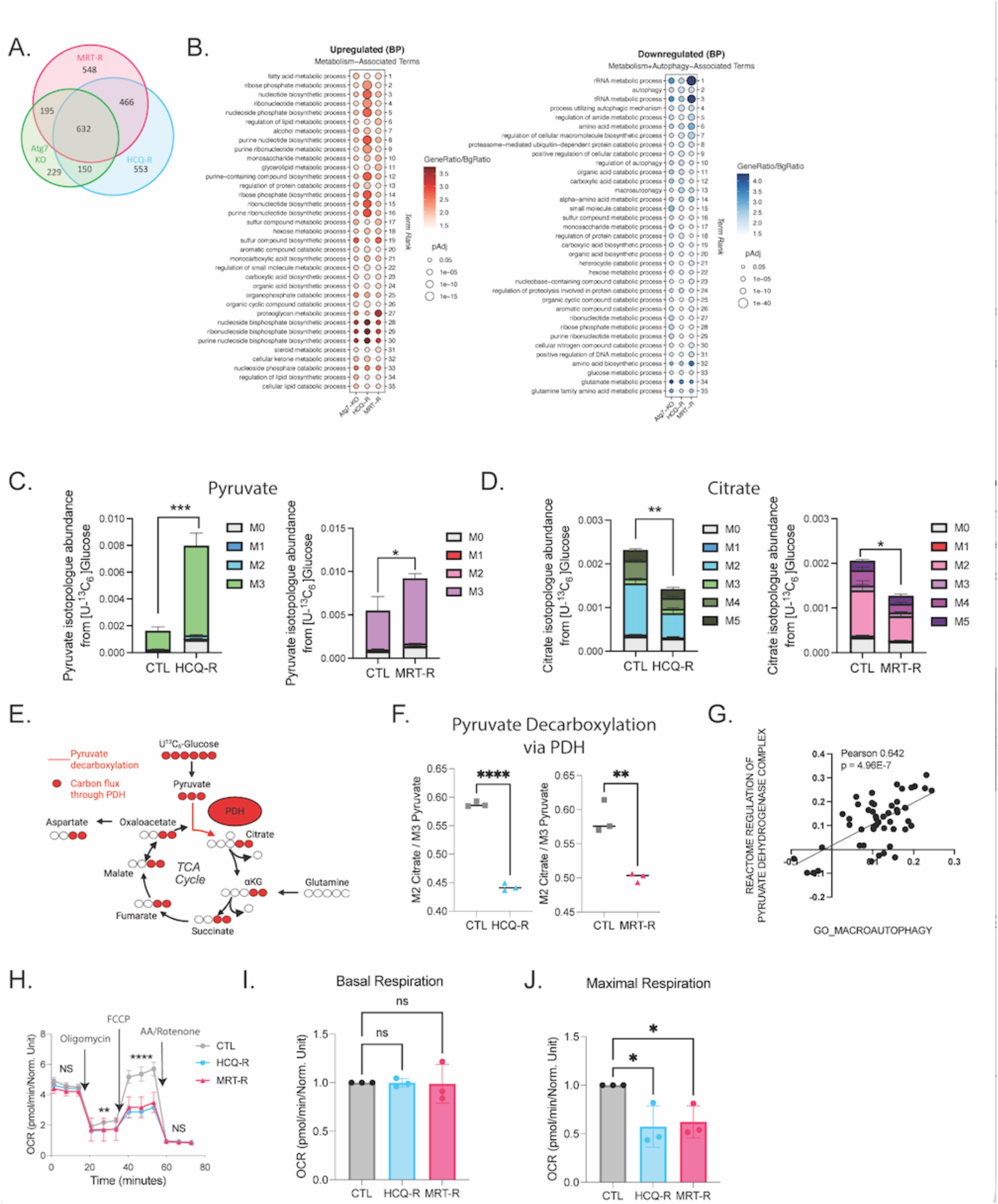
PDA cells resistant to autophagy inhibition rewire their metabolism and decrease pyruvate decarboxylation. (A) Overlapping GO terms from RNA sequencing performed in FC1199 HCQ-R and MRT-R cells compared to CTL, and from FC1199 Atg7 KO compared to NT controls. (B) Top metabolism-related GO terms pathway significantly upregulated (left) or downregulated (right) in FC1199 HCQ-R, MRT-R and Atg7 KO cells. (C-D) Pyruvate (C) and citrate (D) isotopologue abundances in FC1199 CTL, HCQ-R and MRT-R cells after a 24h [U-^13^C_6_] glucose trace. (E) Tracing map for [U-^13^C_6_] glucose trace into pyruvate decarboxylation and TCA cycle. (F) Pyruvate decarboxylation measured as M2 Citrate to M3 pyruvate ratio from the [U-^13^C_6_] glucose trace in FC1199 CTL, HCQ-R and MRT-R cells. (G) Correlation GO macroautophagy gene signature and the reactome regulation of PDH signature in PDA cells determined by single linear regression using Depmap data. (H-J) Seahorse assay performed in FC1199 CTL, HCQ-R and MRT-R cells (H) with calculated basal (I) and maximal (J) respiration rates displayed as the fold change relative to CTL. Data are represented as a mean ± SEM. Data on panels C, D F and H represent n=3-6 technical replicates representative of n=3 biological replicates. Data on panels I and J are representative of n=3 biological replicates. ^∗^ indicates p<0.05, ^∗∗^ p<0.001, ^∗∗∗^ p <0.0001, ^∗∗∗∗^ p<0.00001, determined by two-tailed Student’ *t* test (C-D and F) and one-way ANOVA with Tukey’s post hoc test (H-J). MRT68921 abbreviated as MRT.

Given these alterations in pyruvate metabolism, a process largely isolated to the mitochondria, we assessed mitochondrial function more broadly. Seahorse assays were performed to measure oxygen consumption rates (OCR) as a proxy for oxidative phosphorylation. Mitochondrial stress tests were performed using oligomycin, an ATP synthase (Complex V) inhibitor that stops ATP production followed by the FCCP uncoupler which collapses the proton gradient and maximizes the electron transport chain reaction and antimycin A and rotenone to inhibit Complex I and II and stop all mitochondrial respiration. These assays revealed that the HCQ-R and MRT-R cells have defective mitochondria that cannot reach the same maximal respiration as the CTL cells (Figure 2H-J). Similar results were seen in FC1245 cells with acquired resistance to autophagy inhibition (Figure S3A).

To further characterize TCA function alterations, we performed a [U-^13^C_5_] glutamine trace as glutamine is a key anaplerotic substrate that replenishes TCA cycle intermediates. Glutamine is primarily consumed through its conversion to glutamate followed by either deamination or - transamination to form ⍺-ketoglutarate (⍺KG), which then enters the TCA cycle where it progresses to oxaloacetate and exits as either pyruvate or aspartate. [U-^13^C_5_] glutamine tracing results in M5 ⍺KG and forward cycling in the TCA results in M4 labeling on all TCA intermediates (Figure S3B). We found that glutamine utilization in the TCA cycle was significantly reduced in the HCQ-R and MRT-R cells compared to drug naïve CTL, as we observed decreased M5 ⍺KG, M4 fumarate, M4 malate, M4 aspartate and M4 citrate (Figure S3C-G). These findings together with the [U-^13^C_6_] glucose trace confirm that the TCA function is altered in autophagy inhibition resistant cells as glucose or glutamine are less utilized in the TCA cycle in HCQ-R and MRT-R cells compared to controls.

### PDA cells resistant to autophagy inhibition undergo anaplerosis to limit TCA dysfunctions

Surprisingly, the resistant cells have equal oxygen consumption rates under basal conditions suggesting the mitochondria are relatively functional, in the absence of stress (Figure 2I). This, coupled with the fact that resistant cells are actively proliferating, indicates the cells can circumvent decreased pyruvate decarboxylation to sustain TCA metabolism. Since incorporation of glutamine-derived carbons into the TCA cycle is reduced in autophagy inhibition resistant cells, we asked whether TCA intermediates could be replenished by additional anaplerotic reactions. Reductive carboxylation is a metabolic process enabling the cells to convert ⍺KG into isocitrate, and then to citrate via isocitrate dehydrogenase (IDH), therefore operating in reverse of the TCA direction. In a [U-^13^C_5_] glutamine trace, reductive carboxylation results in M5 citrate, instead of M4 citrate when oxidative TCA metabolism progresses^38^. We also found a modest but significant increase in M5 citrate in HCQ-R cells compared to controls, indicative of a slight increase in reductive carboxylation (Figure S3D).

Another major anaplerotic pathway used to replenish TCA intermediates is pyruvate carboxylation. As previously mentioned, glucose-derived pyruvate can enter the TCA cycle via the PDH-mediated pyruvate decarboxylation, but the other major fate of glycolytic pyruvate is pyruvate carboxylation (PC) via the rate limiting enzyme pyruvate carboxylase (PC). This is an irreversible reaction whereby pyruvate is converted into oxaloacetate. Pyruvate anaplerosis through PC can be assessed by a [U-^13^C_6_] glucose trace, where M3 pyruvate is converted to M3 oxaloacetate via PC resulting in M3 aspartate, M3 fumarate, and M3 malate^39^. However, all of these isotopologues can also be formed via multiple oxidation rounds in the TCA cycle. To accurately measure pyruvate anaplerosis, M3 aspartate, fumarate, and malate can instead be compared to M3 succinate, effectively normalizing pyruvate anaplerosis to multiple oxidation rounds of the TCA cycle^39^ (Figure 3A). We found that HCQ-R and MRT-R cells have an increase in M3 fumarate, M3 malate, and M3 aspartate when compared to M3 succinate indicating increased pyruvate carboxylation compared to controls (Figure 3B-D). To validate this, we used a positional tracer, [3-^13^C_1_] glucose, where only the third carbon of glucose is labelled. [3-^13^C_1_] glucose is converted into [1-^13^C_1_] pyruvate (Figure 3E). With this tracer, the ^13^C is lost in the PDH reaction and only retained through the PC reaction resulting in M1 malate and M1 aspartate. Using this tracer, we found that the autophagy inhibition resistant cells have increased M1 aspartate and M1 malate over M1 pyruvate compared to drug naïve CTL cells (Figure 3F-G). In line with these results, PC is upregulated at both the transcript (*Pcx* gene) and protein level in the HCQ-R and MRT-R cells (Figure 3H-I). Together, these results show that PDA cells that adapt to autophagy inhibition have altered TCA metabolism and glutamine anaplerosis. However, the resistant cells can compensate by increasing reductive carboxylation and PC as a means to replenish TCA intermediates.

**Figure 3.**
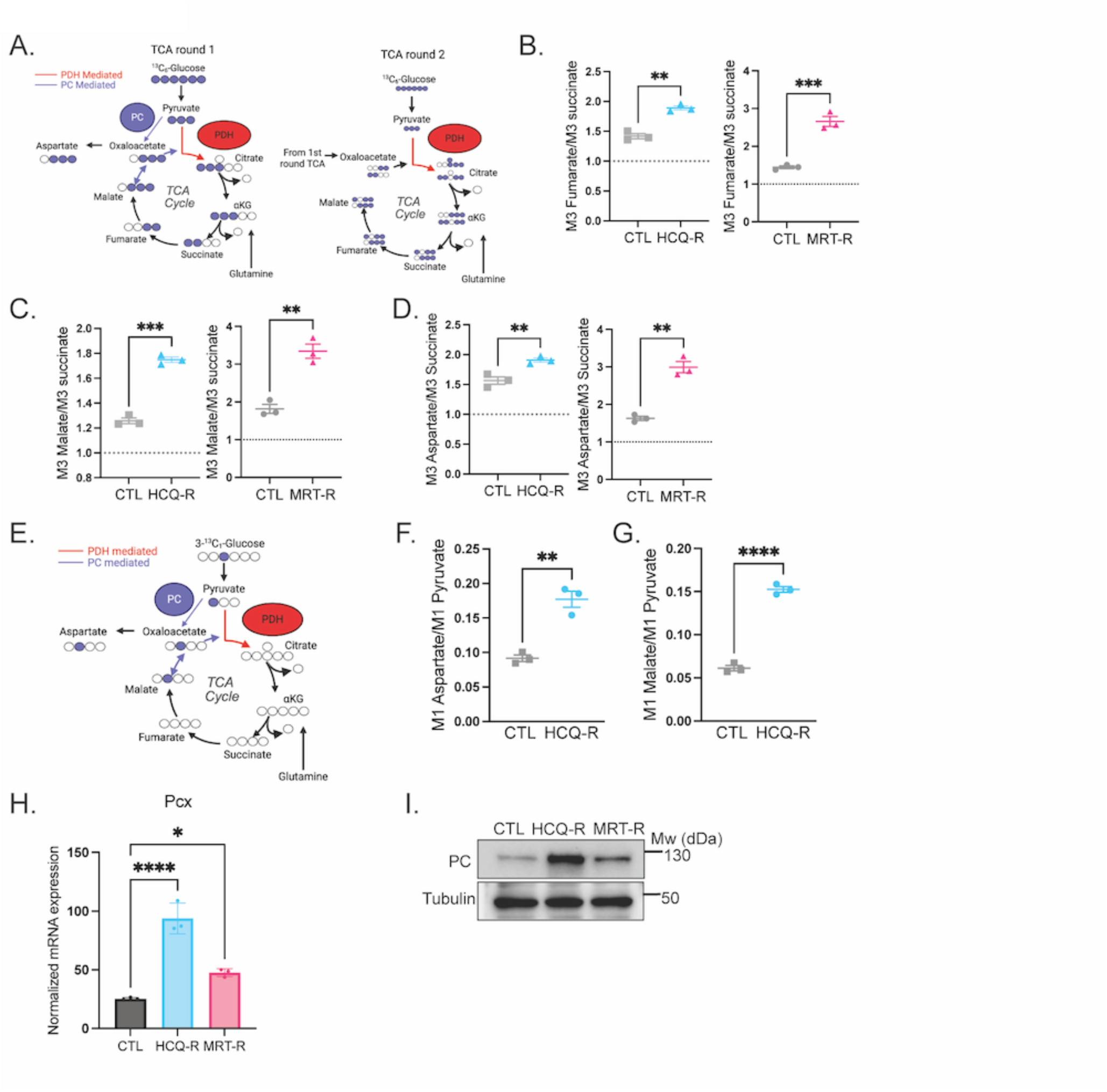
PDA cells resistant to autophagy inhibition undergo anaplerosis to limit TCA dysfunctions. (A) Tracing map for [U-^13^C_6_] glucose trace into pyruvate anaplerosis and TCA cycle round 1 (left) and round 2 (left). (B-D) Pyruvate anaplerosis measured as M3 Fumarate/M3 succinate (B), M3 Malate/M3 succinate (C) or M3 Aspartate/M3 succinate (D) from the [U-^13^C_6_] glucose trace in FC1199 CTL, HCQ-R and MRT-R cells. Reference dotted line at 1.0 represents no pyruvate anaplerosis. (E) Tracing map for [3-^13^C_1_] glucose trace into pyruvate anaplerosis. (F-G) Pyruvate anaplerosis measured as M1 aspartate/M1 pyruvate ratio (F) and M1 malate/M1 pyruvate (G) from the [3-^13^C_1_] glucose trace in FC1199 CTL and HCQ-R cells. (H) *Pcx* normalized mRNA expression in FC1199 CTL, HCQ-R and MRT-R cells. (I) Western blots for PC and Tubulin in FC1199 CTL, HCQ-R and MRT-R cells. Images are representative of n=3 biological experiments. Data are represented as a mean ± SEM. Panels B-D and F-G represent n=3 technical replicates. Panel H represents n=3 biological replicates. ^∗^ indicates p<0.05, ^∗∗^ p<0.001, ^∗∗∗^ p <0.0001, ^∗∗∗∗^ p<0.00001, determined by two-tailed Student’ *t* test (B-D and F-G) and one-way ANOVA with Tukey’s post hoc test (H). MRT68921 abbreviated as MRT.

### PDA cells resistant to autophagy inhibition have decreased nucleotide pools and altered pyrimidine metabolism

Recently, Sahu et al. found that decreased pyruvate decarboxylation can be caused by decreased pyrimidines, one of the two families of nucleotides that make up nucleic acids. UTP is the main phosphodonor for the phosphorylation of thiamine into thiamine pyrophosphate (TPP) by thiamine pyrophosphokinase 1 (TPK1), and TPP positively regulates PDH activity ^40^. Therefore, low UTP causes a decrease in TPK1 activity, effectively reducing PDH-mediated pyruvate decarboxylation. Interestingly, *Tpk1* mRNA expression was significantly decreased in HCQ-R and MRT-R cells compared to drug naïve CTL cells (Figure 4A). Furthermore, gene set enrichment analyses (GSEA) on DEGs from RNA-seq showed that nucleotide metabolism and particularly pyrimidine metabolism-related pathways were significantly altered in HCR and MRT-R cells compared to drug-naïve CTL cells (Figure 4B). We used liquid chromatography mass spectrometry (LC-MS) and performed metabolomics assay to assess free nucleotide pools in the cells with acquired resistance to autophagy inhibition and found that nucleotide pools, including pyrimidine pools, were decreased compared to CTL cells (Figure S4A-B). Cytidine monophosphate (CMP) and uridine monophosphate (UMP) were consistently decreased in HCQ-R and MRT-R cells compared to controls (Figure 4C-D). If the resistant cell lines have lower free pyrimidine pools, but are still actively proliferating, we reasoned that they must utilize pyrimidine metabolism pathways to generate and incorporate pyrimidines into the critical biomass of RNA and DNA.

**Figure 4:**
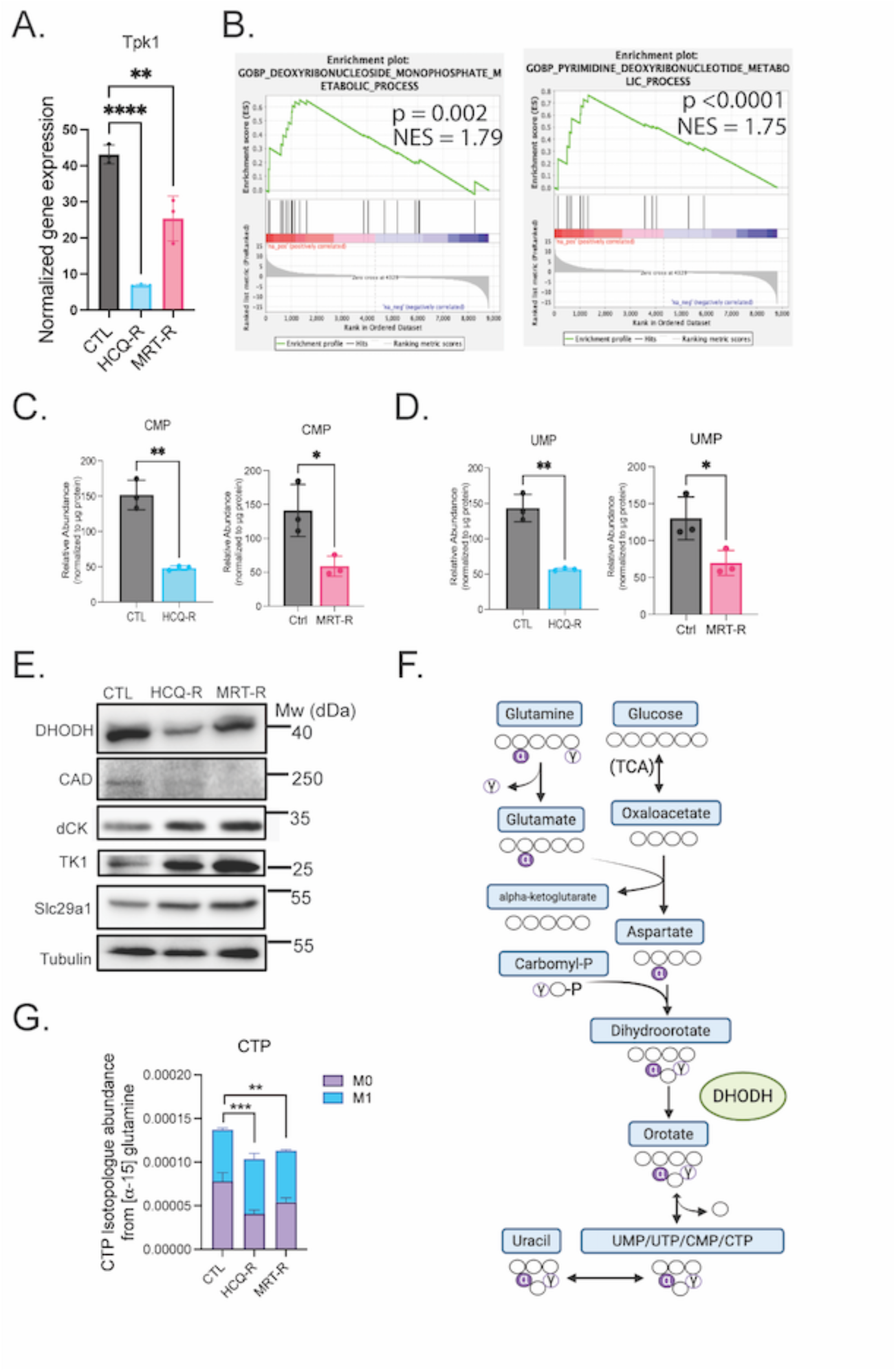
PDA cells resistant to autophagy inhibition have decreased nucleotide pools and altered pyrimidine metabolism. (A) *Tpk1* normalized mRNA expression in FC1199 CTL, HCQ-R and MRT-R cells. (B) GSEA analysis for pyrimidine metabolism related processes in FC1199 CTL and HCQ-R cells. (C-D) CMP (D) and UMP (D) levels in FC1199 CTL, HCQ-R and MRT-R cells. (E) Western blot for DHODH, CAD, DCK, TK1, SLC29A1 and Tubulin in FC1199 CTL, HCQ-R and MRT-R cells. Images are representative of n=3 biological experiments. (F) Tracing map for [⍺-^15^N] glutamine trace into de novo pyrimidine synthesis. (G) CTP isotopologue abundance in FC1199 CTL, HCQ-R and MRT-R cells after a 24h [⍺-^15^N] glutamine trace. Data are represented as a mean ± SEM. Panels A and C-D represent n=3 biological replicates. Panel G represents n=3 technical replicates. ^∗^ indicates p<0.05, ^∗∗^ p<0.001, ^∗∗∗^ p <0.0001, ^∗∗∗∗^ p<0.00001, determined by one-way ANOVA with Tukey’s post hoc test (A and G) and two-tailed Student’ *t* test (C-D). MRT68921 abbreviated as MRT.

Most cells can synthesize pyrimidines from aspartate and also salvage pyrimidines. The rate limiting enzymes for the de novo pyrimidine pathway are Carbamoyl-phosphate synthetase 2, Aspartate transcarbamoylase, and Dihydroorotase (CAD) and dihydroorotate dehydrogenase (DHODH). CAD catalyzes the first three steps of this reaction, transforming glutamine, aspartate, and bicarbonate into dihydroorotate through a series of energy-consuming reactions, and DHODH converts dihydroorotate into orotate which is subsequently converted to UMP and all other pyrimidines with phosphate esters^41,42^. On the other hand pyrimidine salvage recovers extracellular nucleosides and nucleobases from the extracellular environment or from intracellular nucleic acid degradation to synthesize nucleotides^41,42^. Extracellular pyrimidines are transported into the cell through ENT nucleoside transporters encoded by *Slc29a1-4*. Salvaged nucleosides are converted into their corresponding nucleoside monophosphates through phosphorylation reactions^41,43^. For example, deoxycytidine requires deoxycytidine kinase (dCK) to form deoxycytidine monophosphate (dCMP), thymidine uses thymidine kinase (TK1) to form deoxythymidine monophosphate (dTMP).

We found that the HCQ-R and MRT-R resistant cell lines have decreased expression in the rate-limiting enzymes for the de novo synthesis pathway but increased expression in pyrimidine salvage enzymes and transporters (Figure 4E, S4C). Next, we performed an [⍺-^15^N] glutamine trace where the ⍺-^15^N is metabolized through aspartate onto dihydroorotate and orotate via DHODH, and subsequently incorporated into pyrimidines through UMP to CMP and CTP (Figure 4F). A 24hr trace with an [⍺-^15^N] glutamine confirmed that the resistant cells do not upregulate de novo pyrimidine metabolism (Figure 4G). Without an increase in de novo pyrimidine metabolism, we reasoned that the cells resistant to autophagy inhibition must increase pyrimidine salvage in attempts to mitigate their decreased pyrimidine pools to sustain active proliferation and biosynthesis.

### PDA cells resistant to autophagy inhibition have increased sensitivity to pyrimidine analogs

Considering the perturbations observed in pyrimidine metabolism in the HCQ-R and MRT-R cell lines, most notably upregulation of the pyrimidine salvage pathway, we hypothesized that the resistant cells would also increase the salvage of pyrimidine analogues through this same pathway. Indeed, prior studies found that *dCK* expression correlates with sensitivity to the deoxycytidine analogue, gemcitabine (2′,2′-difluorodeoxycytidine) ^44–47^. Similarly, *TK1* expression correlates with sensitivity to the thymidine analogue, trifluridine (FTD: 5-trifluromethyl-2’-deoxyuridine)^48,49^. Trifluridine is most effective when combined with the thymidine phosphorylase inhibitor, tipiracil (TPI) to prevent its rapid degradation and is administered as FTD/TPI ^54,5^.

To test sensitivity to pyrimidine analogues, we performed cell viability assays in the HCQ-R, MRT-R, and CTL cells treated with a panel of drugs including current PDA standard of care, and targeted agents tested in clinical trials, including pyrimidine analogs. The FC1199 HCQ-R and MRT-R cells displayed near pan-resistance to all compounds except for the pyrimidine analogs gemcitabine, FTD/TPI, and cytarabine (another cytidine analogue with a modified sugar moiety; 1-.beta.-D-arabinofuranosyl-4-amino-2(1H)pyrimidinone) (Figure 5A-B). Interestingly, fluorouracil (5-FU; 5-fluoro-2,4-(1H,3H)-pyrimidinedione) is also considered a pyrimidine analogue, however it is metabolized through the de novo pyrimidine synthesis pathway^50,51^, unlike gemcitabine, FTD/TPI, and cytarabine – all of which are which are metabolized through the pyrimidine salvage pathway. Accordingly, the HCQ-R and MRT-R cells are not more sensitive to 5-FU (Figure 5A). Moreover, the cells resistant to autophagy inhibition are not more sensitive to other small molecules that target de novo pyrimidine metabolism like Brequinar or BAY-2402234, both of which target DHODH (Figure S4D-G).

**Figure 5.**
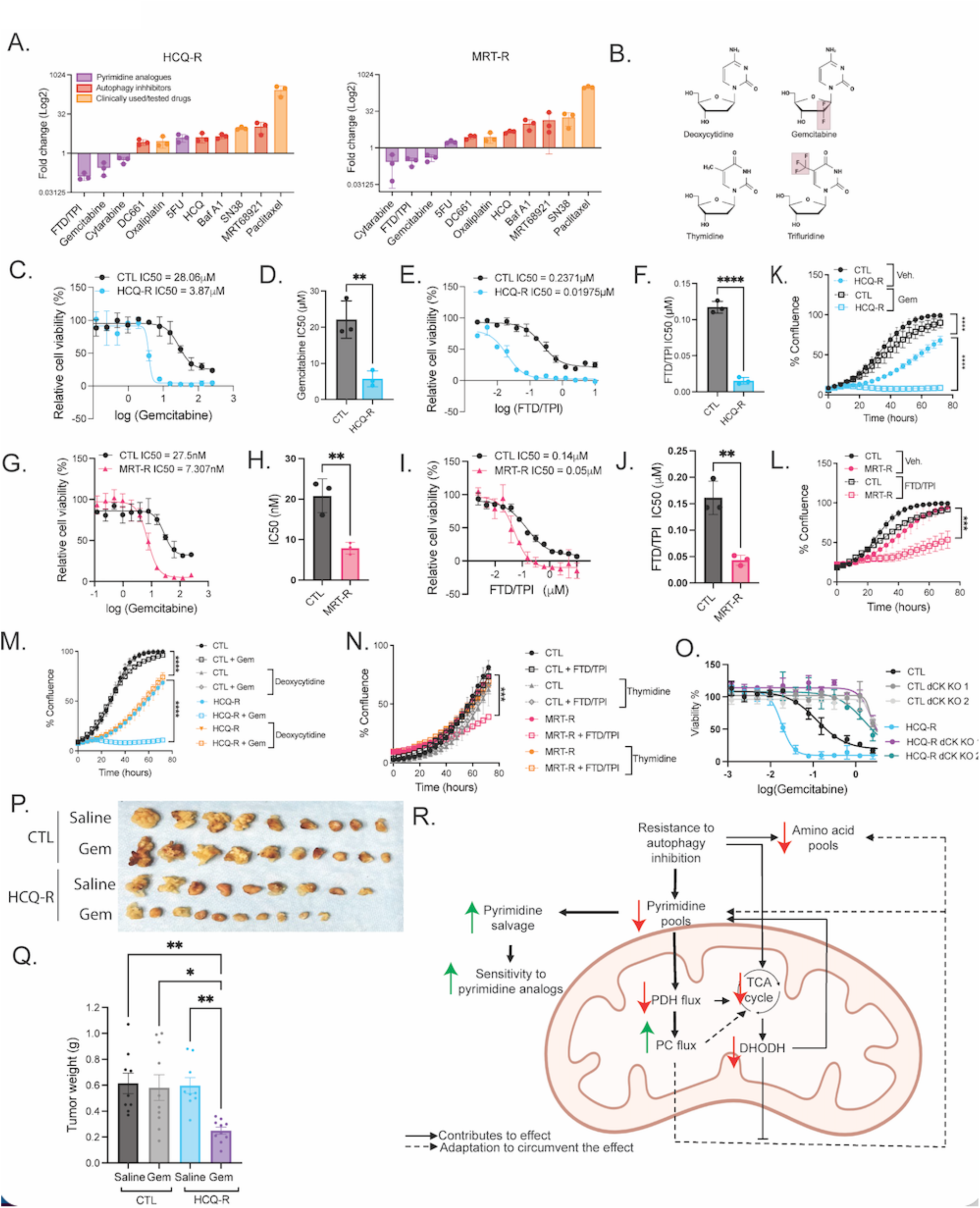
PDA cells resistant to autophagy inhibition have increased sensitivity to pyrimidine analogs. (A) Log2 fold change between the IC50 of FC1199 HCQ-R and CTL cells or FC1199 MRT-R and CTL cells for the indicated autophagy inhibitors, pyrimidine analogs and other PDAC standard of care determined with MTT cell viability assays. (B) Structure of deoxycytidine compared to gemcitabine and thymidine compared to trifluridine. (C-F) MTT cell viability assay and the calculated IC50 for FC1199 CTL and HCQ-R cells treated with Gemcitabine (C-D) and FTD/TPI (E-F) (G-J) MTT cell viability assay and the calculated IC50 for FC1199 CTL and MRT-R cells treated with Gemcitabine (G-H) and FTD/TPI (I-J) (K-L) Incucyte live cell imaging in FC1199 CTL and HCQ-R cells treated with gemcitabine (K) or FC1199 CTL and MRT-R cells treated with FTD/TPI (L) as indicated for 3 days. (M)) Incucyte live cell imaging in FC1199 CTL and HCQ-R cells treated with gemcitabine or vehicle control with or without the addition of deoxycytidine as indicated for 3 days. (N) Incucyte live cell imaging in FC1199 CTL and MRT-R cells treated with FTD/TPI or vehicle control with or without the addition of thymidine as indicated for 3 days. (O) MTT cell viability assay in FC1199 CTL (NT or DCK KO) and HCQ-R cells (NT or DCK KO) treated with gemcitabine for 72h. (P-Q) Picture (P) and weight (Q) of the orthotopic tumors resected from mice injected with FC1245 CTL or HCQ-R cells treated with gemcitabine or saline control (n=9-10/group). (R) Schematic overview of the mechanisms identified or hypothesized in PDA cells resistant to autophagy inhibitors. Data are represented as a mean ± SEM. Panels A, D, F, H, and J represent n=3 biological replicates. Panel D-G represents n=3 technical replicates representative of n=3 biological replicates. ^∗^ indicates p<0.05, ^∗∗^ p<0.001, ^∗∗∗^ p <0.0001, ^∗∗∗∗^ p<0.00001, determined by two-tailed Student’ *t* test (A, D, F, H, and J), two-way repeated measures ANOVA with Tukey’s post hoc test (K-N) and one-way ANOVA with Tukey’s post hoc test (Q). MRT68921 abbreviated as MRT.

Gemcitabine and FTD/TPI are of particular interest because are in widespread clinical use. Gemcitabine is an FDA standard-of-care agent for PDA treatment and FTD/TPI is used for colorectal and gastric cancers^52–54^. FTD/DTPI is currently being tested in clinical trials for PDA patients in combination with paclitaxel (NCT04046887). The HCQ-R and MRT-R cells have a 3-5-fold decrease in the IC50 for gemcitabine and FTD/TPI (Figure 5C-J). Similarly, the FC1245 and Panc1 cells with acquired resistance to autophagy inhibition are also hypersensitive to these pyrimidine analogues (Figure S5A-F). Incucyte live-cell imaging confirmed that low doses of gemcitabine and FTD/TPI, that have minimal effects in the CTL cells, drastically limit cell growth in the HCQ-R and MRT-R lines (Figure 5K-L, S5G).

Gemcitabine incorporation into DNA interferes with DNA synthesis and inhibits cell proliferation. Gemcitabine also inhibits ribonucleotide reductase (RNR), which catalyzes the formation of deoxyribonucleotides ^55,56^. To assess if gemcitabine sensitivity is due to increased salvage because it resembles a pyrimidine, we tested the effects of another small molecule, triapine – a thiosemicarbazone, that also targets RNR, but is not a pyrimidine analogue^57^. Interestingly, the FC1199 and FC1245 HCQ-R and MRT-R cells are resistant to triapine, showing opposite effects to what was seen with gemcitabine (Figure S5H-K). The pyrimidine structure of gemcitabine is likely necessary for its increased uptake in the autophagy-inhibitor-resistant cells. To further explore if the increased sensitivity to these compounds is due to increased salvage, we tested if the addition of extracellular pyrimidines could rescue these effects. The addition of deoxycytidine or uridine abrogated the increased sensitivity to gemcitabine in the FC1199 and FC1245 HCQ-R cells (Figure 5M, S5L-M). Similar effects were found in FC1199 MRT-R cells treated with FTD/TPI, and the effects could be rescued with the addition of either thymidine or uridine (Figure 5N, S5N). Interestingly, the fact that uridine phenocopies cytidine in these assays indicates that the low intracellular pyrimidine pools are leading to a broad upregulation of pyrimidine salvage. Uridine can restore CMP and TMP levels in cells – since uridine-derived UMP is readily converted to these other pyrimidines – thereby rescuing sensitivity to pyrimidine analogues. Genetic knock out of dCK, required to metabolize salvaged pyrimidines as well as gemcitabine, eliminated any differential sensitivity between the HCQ-R and CTL cells (Figure 5O, S5O). Together these studies demonstrate that cells that acquire resistance to autophagy inhibition, upregulate pyrimidine salvage, and consequently increase their uptake pyrimidine analogues leading to hypersensitivity.

To determine if this acquired sensitivity to pyrimidine analogues is maintained in vivo we orthotopically transplanted the HCQ-R and CTL cells into the pancreas of C57Bl/6 mice. Prior to this, we first confirmed that the autophagy-inhibitor-resistant cell lines maintain sensitivity to gemcitabine even if they are grown without continuous dosing of the autophagy inhibitors. Cell viability assays showed that they retained their resistance to autophagy inhibitors as well as their increased sensitivity to gemcitabine (Figure S5P-U). This allows us to remove the cells from drug to transplant them in vivo. We injected the FC1245 and FC1199 HCQ-R cells orthotopically in the pancreas and found that a low gemcitabine dose did not impact CTL tumor growth but significantly reduced HCQ-R tumor volume, despite equal tumor size prior to drug treatment (Figure 5P-Q, S5V-W).

All together these studies lead us to propose a mechanistic model stating that as cells acquire resistance to autophagy inhibition, they rewire their metabolism. This results in an overall decrease in pyrimidine pools as well as altered TCA metabolism including decreased pyruvate decarboxylation. As a work-around, the resistant cells increase pyruvate anaplerosis to replenish TCA intermediates and generate aspartate to replenish pyrimidine pools. However, since DHODH and de novo pyrimidine metabolism do not also increase to match this need, the cells must salvage pyrimidines from the extracellular environment. This adaptation leads to increased pyrimidine salvage and causes the cells to increase salvage of pyrimidine analogues, resulting in a hypersensitivity to different pyrimidine analogues (Figure 5R).

### Autophagy inhibitors and pyrimidine analogues have a combinatory effect in PDA

These new mechanistic insights led us to test the effects of combining autophagy inhibition and pyrimidine analogues. We used a biobank of pharmacotyped PDA patient-derived organoids (PDOs). Human-derived PDA organoids are a translationally relevant model that can predict response to treatment in patients ^58^. We compared RNA-seq and sensitivity to gemcitabine in a cohort of 43 PDOs. The original tumor samples were obtained from patients of both sexes (20 women, 23 men), and the majority were derived from the primary PDA, with six derived from metastatic sites and two from ascites. 40 PDO’s have *KRAS* mutations (including G12D, G12V, G12R and Q61H) while three maintain WT *KRAS*. We found a significant correlation between the autophagy-lysosome gene signature^37^ and gemcitabine sensitivity across PDOs showing that low autophagy gene expression correlates with higher gemcitabine sensitivity (Figure 6A, S6A).

**Figure 6:**
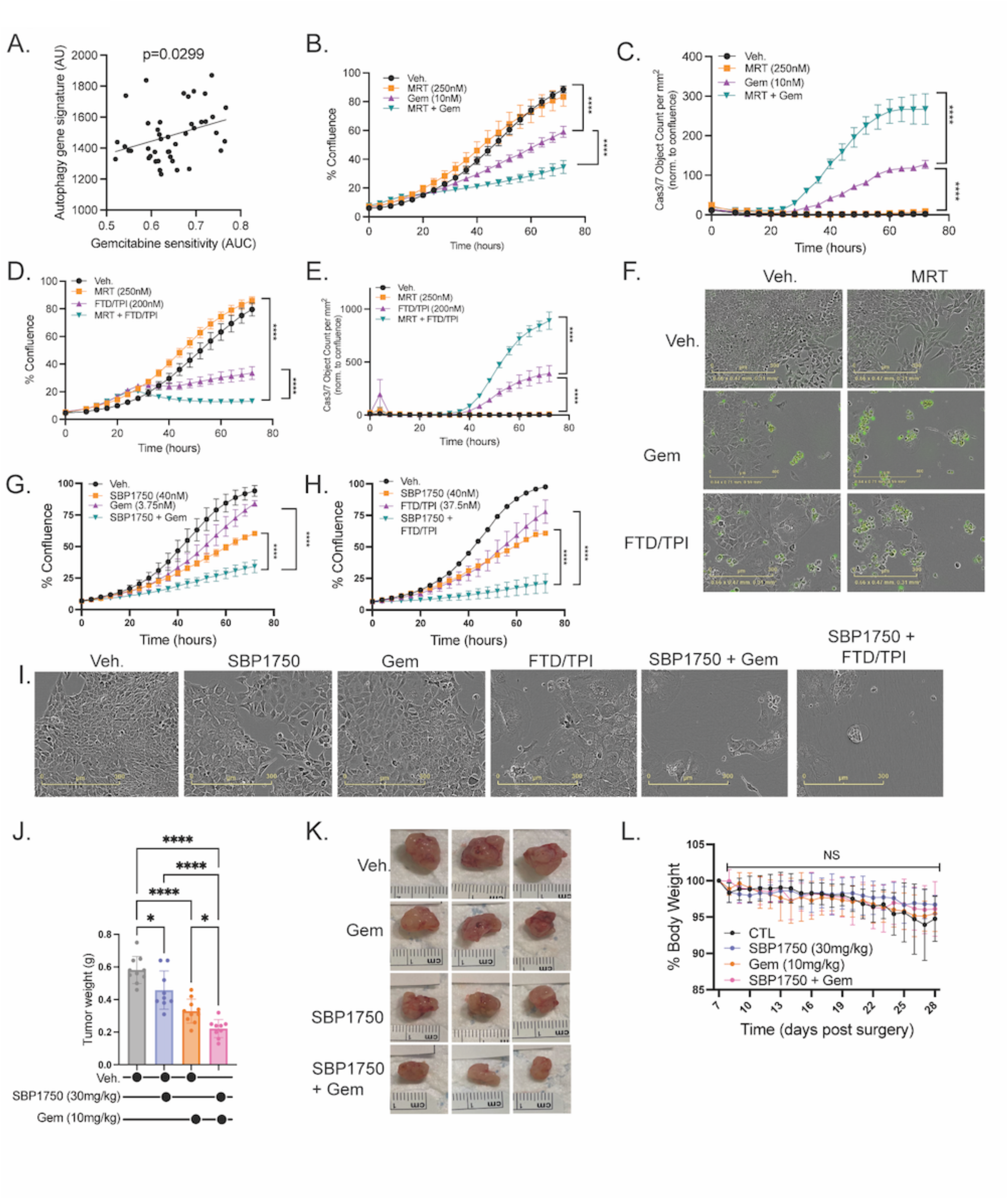
Autophagy inhibitors and pyrimidine analogues have a combinatory effect in PDA. (A) Correlation between autophagy gene signature and gemcitabine sensitivity in hPDOs determined by single linear regression. (B-I) Incucyte live cell imaging in FC1199 cells treated as indicated for 3 days and graphed as percent confluence (B,D,G,H) or caspase 3/7 object count/mm^2^ normalized to confluence (E,H) with (F) showing representative phase imaging and cas3/7 object count from (C and E) and (I) showing representative phase imaging from (G and H). (J-L) Tumor volumes (J) from mice injected orthotopically with FC1199 cells and treated with gemcitabine and/or SBP-1750 or vehicle control with (K) showing representative images of tumors and (L) changes in body weight starting 7 post orthotopic injection on day 1 of treatment (K). n = 9-10 per group. Data are represented as a mean ± SEM. Data on panels B-E and G-H represent n=3 technical replicates representative of n=3 biological replicates. ^∗^ indicates p<0.05, ^∗∗^ p<0.001, ^∗∗∗^ p <0.0001, ^∗∗∗∗^ p<0.00001, determined by two-way repeated measures ANOVA with Tukey’s post hoc test (B-E, G-H and L) and one-way ANOVA with Tukey’s post hoc test (J). MRT68921 abbreviated as MRT.

We assessed the effects of combining autophagy inhibitors HCQ or MRT68921 with gemcitabine or FTD/TPI in FC1199, FC1245 and Panc1 drug naïve cells. Incucyte live-cell imaging and clonogenic growth assays revealed that all combinations with autophagy inhibitors and pyrimidine analogues resulted in a greater effect on cell viability than any single agent (Figure 6B-F, S6B-K). HCQ has poor bioavailability and specificity, while targeting ULK1/2 kinases may overcome these limitations, representing a more attractive therapeutic strategy. Importantly, MRT68921 showed a robust combinatory effect with both gemcitabine and FTD/TPI resulting in decreased cell survival and increased apoptosis, as measured by cleaved caspase activity with Incucyte live cell imaging (Figure 5B-F, S6G-J). Despite these promising in vitro studies with MRT68921, this drug showed no single agent effect in vivo suggesting it too has poor bioavailability.

Shaw et al. have recently designed and synthesized potent small molecule inhibitors of the ULK1/2 kinases, including SBP-1750^59^. This compound possesses good oral bioavailability and exposure, highlighting its potential as a promising candidate for combination therapy studies. Given this, we combined SBP-1750 with pyrimidine analogues and found a robust combinatory effect with both gemcitabine and FTD/TPI (Figure 6G-I). To confirm this effect in vivo, we orthotopically transplanted FC1199 cells into the pancreas of C57Bl/6 mice and found that combined treatment with SBP-1750 and a low dose of gemcitabine resulted in the most robust decrease in tumor size (Figure 6J-K). Importantly, this new therapeutic strategy did not lead to dose-limiting toxicities (Figure 6L). Together these studies indicate that autophagy inhibition causes altered pyrimidine metabolism which can be leveraged to increase sensitivity to clinically relevant pyrimidine analogues like gemcitabine.

## DISCUSSION

Numerous in vitro and physiologically relevant in vivo studies confirm that autophagy is a valid target to treat pancreatic cancer ^9,22,10,60,23,16^. However, the field lacked a mechanistic understanding of how autophagy inhibition rewires cell metabolism, resulting in ineffective therapeutic strategies that lead to acquired resistance. In this study, we generated PDA cells resistant to autophagy inhibitors to better understand these acquired adaptations and in the process, we identified unique metabolic vulnerabilities.

For the first time, we found that that PDA cells can adapt to the lysosomal inhibitor HCQ and to the ULK1/2 inhibitor MRT68921 (Figure 1). These two drugs have very different mechanisms of action – with one targeting downstream lysosomes and the other targeting upstream kinases – but bulk RNA-seq revealed that PDA cells resistant to HCQ and MRT68921 share numerous adaptative pathways compared to drug naïve control cells. Therefore, many of these shared pathways are likely specific to autophagy inhibition rather than representing off-target effects. Notably, multiple metabolism-related processes were identified. This is unsurprising as autophagy participates in maintaining metabolic homeostasis in the cells by degrading proteins, organelles, and cytoplasmic components to generate substrates such as amino acids, fatty acids, sugars, and nucleosides ^61,62^. These critical building blocks and metabolites can further fuel key metabolic pathways and biosynthesis. Ubiquitous loss of key autophagy genes like *Atg5* or *Atg7* leads to severe metabolic dysfunction, including energy depletion, impaired lipid and glycogen mobilization, and ultimately early death due to metabolic insufficiency ^13,63,64^.

We found that autophagy-inhibition-resistant cells cannot perform autophagy but can still continue to proliferate, suggesting a reprogramming of cell metabolism (Figure 1, S1). With metabolomics assay, we found an accumulation of pyruvate in the HCQ-R and MRT-R cells compared to drug naïve cells. Stable isotope tracing of glucose and glutamine revealed that pyruvate metabolism was rewired in cells resistant to autophagy inhibitors and pyruvate entry into the TCA cycle via PDH was decreased (Figure 2, S2, S3). Even though PDH links glycolysis to the TCA function, basal respiration rates were surprisingly not impacted, and only maximal respiration rates were significantly lower in the resistant cells compared to controls. We found that HCQ-R and MRT-R cells had increased pyruvate anaplerosis via pyruvate carboxylation, allowing the resistant cells to replenish TCA intermediates and compensate, at least partly, for decreased PDH activity (Figure 3). These results aligns with previous studies in non-cancer cells with dysfunctional PDH activity (*Pdha1* KO) which also found an increase in PC activity ^65,66^, and for the first time link autophagy to pyruvate carboxylation and decarboxylation.

We speculate that the link between autophagy and decreased PDH-mediated pyruvate decarboxylation is pyrimidine pool levels. A recent report found that PDH activity is dependent on pyrimidine levels, and UTP is used as the phospho-donor by TPK1 to phosphorylate thiamine into TPP, which in turn activates PDH^40^. Gene expression changes indicate decreased *Tpk1* expression as well as altered pyrimidine metabolism in the autophagy-inhibitor-resistant cells. We confirmed decreased nucleotide pools, including pyrimidine pools, in HCQ-R and MRT-R cells compared to controls (Figure 4, S4). This supports previous findings linking autophagy to nucleotide metabolism: in murine lung cancer cells lacking *Atg7*, it was shown that nucleotide pools were significantly decreased under starvation conditions compared to WT control cells^67^. Furthermore, short-term chloroquine treatment in PDA cells has also been shown to reduce nucleotide pools^68^.

Most importantly, despite a decrease in nucleotide pools, the autophagy-deficient cells are still actively proliferating, and therefore must find ways to replenish the nucleotides required for biosynthesis of DNA and RNA. In addition to recycling nucleic acids to replenish pyrimidine pools, cells can also de novo synthesize pyrimidines as well as salvage pyrimidines from the extracellular environment. In exploring these two pyrimidine metabolism pathways, we found that as cells adapt to loss of autophagy, they preferentially salvage pyrimidines instead of synthesizing them de novo. De novo synthesis requires the DHODH enzyme, which resides in the mitochondrial inner membrane and its activity is coupled with the electron transport chain. Autophagy is critical for maintaining mitochondrial quality control via mitophagy and loss of autophagy results in the accumulation of damaged mitochondria which leads to altered cell metabolism. Accordingly, DHODH levels were decreased in the HCQ-R and MRT-R making it impossible for the cells to upregulate de novo pyrimidine metabolism to overcome decreased nucleotide pools and sustain biosynthesis (Figure 4).

In contrast, we found that as cells adapt to autophagy inhibition, they preferentially salvage pyrimidines (Figure 4, S4). Importantly, this pathway is not only involved in the salvaging of pyrimidines, but also in the metabolism of pyrimidine analogs. Therefore, instead of trying to thwart resistance to autophagy inhibition, we propose to leverage it. Pyrimidine analogs (e.g., gemcitabine) are commonly used to treat PDA patients but typically show limited efficacy. Our studies show that HCQ-R and MRT-R cells can utilize increased pyrimidine salvage to uptake and metabolize pyrimidine analogs resulting in increased sensitivity to these compounds (Figure 5, S5). We found that autophagy-inhibition-resistant cells are more sensitive to the thymidine analog FTD/TPI, as well as the cytidine analogues gemcitabine and cytarabine. This is in stark contrast to the pan-resistance observed to all autophagy inhibitors and other PDA standard of care. 5-FU was the only pyrimidine analog tested for which the autophagy inhibition resistant cells showed resistance. Interestingly, this compound is predominantly converted to an active anticancer metabolite via Uridine Monophosphate Synthetase (UMPS)^50,51^, which is part of the de novo pyrimidine synthesis pathway rather than the salvage pathway. We confirmed that increased pyrimidine salvage is the mechanism leading to sensitivity to pyrimidine analogs in HCQ-R and MRT-R cells by adding extracellular pyrimidines or by knocking out the critical salvage enzyme, dCK. Furthermore, the increased sensitivity to gemcitabine translated in vivo as orthotopic injections of FC1199 or FC1245 HCQ-R and CTL cells showed that HCQ-R tumors were hypersensitive to an extraordinarily low dose of gemcitabine as compared to CTL tumors.

Through in-depth metabolic characterization of PDA cells with acquired resistance to the autophagy inhibitors, HCQ and MRT68921, we derive a mechanistic model linking autophagy, pyruvate metabolism and nucleotide metabolism (Figure 5R). Autophagy deficiency has several consequences including decreased pyrimidine pools due to a loss of nucleic acid recycling. A further consequence of low pyrimidine pools is decreased PDH activity resulting in decreased pyruvate decarboxylation. Loss of autophagy also leads to mitochondrial dysfunction, evident in our studies by decreased utilization of glucose and glutamine for TCA metabolism. A further consequence of altered mitochondrial function is less DHODH. As the cells attempt to adapt to these metabolic changes, they upregulate pyruvate anaplerosis via pyruvate carboxylation (PC) to replenish TCA intermediates. We postulate that as the cells sense the lower pyrimidine pools, one key reason the cells increase PC is to generate more aspartate that can feed into de novo pyrimidine synthesis. However, because autophagy also decreases mitochondrial function and DHODH, the autophagy-inhibitor-resistant cells cannot use de novo synthesis, which ultimately leads to an accumulation of aspartate and the pyrimidine pools remain low. But cells resistant to autophagy inhibition are still actively proliferating and synthesizing RNA and DNA. To this end, they upregulate pyrimidine salvage to supply the required pyrimidines necessary for biosynthesis and survival without replenishing the pyrimidine pool.

Autophagy inhibition-induced upregulation of nucleotide salvage and pyrimidine analog uptake may therefore represent a valuable therapeutic leverage for PDA patients. We found a positive correlation between autophagy gene signatures and gemcitabine sensitivity in human-derived PDOs. Therefore, low autophagy correlates with increased sensitivity to gemcitabine in this dataset. We found that combining autophagy inhibitors, HCQ or MRT68921, with pyrimidine analogues, gemcitabine or FTD/TPI, resulted in a combinatory cell killing effect (Figure 6). Previously, HCQ and gemcitabine were clinically tested in a Phase II randomized clinical trial in PDA patients^28^ (NCT01506973). While the results showed an increase in response rate with HCQ, there was no improvement in the primary end point of overall survival at 12 months. This result highlights that we need better autophagy inhibitors to combine with pyrimidine analogues. Our mechanistic studies compare HCQ and ULK1/2 inhibition allowing us to draw the important conclusion that autophagy inhibition, irrespective of where in the pathway, alters pyrimidine metabolism. But HCQ is unlikely to be an effective therapeutic drug in the clinic due to its low specificity and bioavailability.

More specific autophagy inhibitors, and in particular more potent ULK1/2 inhibitors, are being developed^69^ ^70^. Targeting ULK1/2 is effective at reducing PDA cell growth^71^. SBP-1750, a newly developed ULK1/2 inhibitor from Shaw et al., exhibits improved bioavailability in mice and oral exposure compared to previously developed ULK1/2 autophagy inhibitors, making it an ideal candidate for our combinatorial studies. We combined SBP1750 with gemcitabine or FTD/TPI and found a robust combinatory effect on PDA cell viability in vitro and tumor growth in vivo (Figure 6). This result emphasizes that a promising new combinatory strategy could be to combine ULK1/2 inhibitors with pyrimidine analogues. Our mechanistic studies show that adaptations to autophagy inhibition led to the most robust sensitivity to pyrimidine analogues, even more so than combining the two drugs in treatment naïve cells, suggest that treating with an autophagy inhibitor prior to a pyrimidine analogue may be the most effective strategy. Modifying the sequence of treatment in this way would allow autophagy inhibition the necessary time to rewire pyruvate and pyrimidine metabolism to increase pyrimidine salvage. Subsequent treatment with pyrimidine analogues could then be more effective as uptake is increased. We propose that pre-clinical and eventually clinical testing of autophagy inhibition could improve pancreatic cancer patient outcomes.

## Supporting information

Key Resource Table

## ACKNOWLEDGMENTS

This work was supported by an NIH grant R00CA245187 (C.G.T), a Pew-Stewart Scholar award 00036071 (C.G.T), a V Foundation scholar award V2023-006 (C.G.T), and a Lustgarten Foundation-AACR Career Development Award for Pancreatic Cancer Research in honor of John Robert Lewis 24-20-67-TOWE (C.G.T), and the Salk Alumni Pos-Doctoral Fellowship (S.D.). The targeted metabolomics on nucleotide pool were supported by the Mass Spectrometry Core of the Salk Institute (RRID:SCR_014843) with funding from NIH-NCI CCSG P30 CA014195, NIH-NIA San Diego Nathan Shock Center P30 AG068635, an NIH S10 award for metabolic instrumentation: S10 OD021815, and the Helmsley Center for Genomic Medicine. RNA sequencing experiments were supported by The Razavi Newman Integrative Genomics and Bioinformatics Core Facility of the Salk Institute (RRID:SCR_014842 and SCR_014846) with funding from NIH-NCI CCSG P30 CA014195, NIH-NIA San Diego Nathan Shock Center P30 AG068635, the NIH-NIA Liver Cancer P01 AG073084-04, the Howard and Maryam Newman Family Foundation and the Helmsley Trust.

## DECLARATIONS

The authors have no competing interests to declare

---

**Figure S1. Various PDA cell lines can acquire resistance to pharmacological inhibition of autophagy, related to Figure 1**

(A-B) MTT cell viability assay in FC1245 CTL and HCQ-R cells treated with HCQ for 72h (A) and calculated IC50 values (B).

(C) Incucyte live cell imaging in FC1245 CTL and HCQ-R cells treated with HCQ as indicated for 3 days.

(D) Clonogenics assay with FC1199 CTL, HCQ-R and MRT-R cells treated with HCQ or MRT68921 as indicated.

(E-G) MTT cell viability assay in Panc-1 CTL and HCQ-R cells treated with HCQ (E) or MRT68921 (F) for 72h and calculated IC50 values for MTT assay (G).

(F) MTT cell viability assay in FC1245 CTL and HCQ-R cells treated with MRT68921 (0.024-100μM) for 72h to determine the IC50 for this drug. Data shown as n=3 technical replicates and is representative of at least 3 biological experiments.

(G) IC50 values of FC1245 CTL and HCQ-R cells treated with MRT68921 for 72h. Data are represented as a mean ± SD, n = 3 biological replicates; two-tailed Student’ *t* test

Data are represented as a mean ± SEM. Data on panels A, C and E-F represent n=3 technical replicates representative of n=3 biological replicates. Data on panels B and G represent n=3 biological replicates. Data on panel D is representative of n=3 biological replicates. ^∗^ indicates p<0.05, ^∗∗^ p<0.001, ^∗∗∗^ p <0.0001, ^∗∗∗∗^ p<0.00001 determined by two-tailed Student’ *t* test (B and G) and two-way repeated measures ANOVA with Tukey’s post hoc test (C). MRT68921 abbreviated as MRT.

**Figure S2: Metabolomics shows reprogramming of pyruvate metabolism in autophagy inhibition resistant cells, related to Figure 2**.

(A) Western blot for ATG7, LC3I/II and β-actin NT control FC1199 ATG7 KO clones. Images are representative of n=3 biological experiments.

(B) Overlapping GO terms upregulated (left) and downregulated (right) in FC1199 HCQ-R and MRT-R compared to CTL and in Atg7 KO cells compared to NT controls.

(C) Metabolomics assay in CTL and HCQ-R cells (left) and in CTL and MRT-R cells (right).

(D-G) Individual MIDs for pyruvate (D-E) and citrate (F-G) in FC1199 CTL, HCQ-R and MRT-R cells following a 24h [U-^13^C_6_] glucose trace.

(H-K) Normalized abundance and individual MIDs for aspartate (H-I) and ⍺KG (J-K) in FC1199 CTL, HCQ-R and MRT-R cells following a 24h [U-^13^C_6_] glucose trace.

Data are represented as a mean ± SEM. Data on panel C and D-K represent n=3 biological replicates and n=3 technical replicates, respectively. ^∗^ indicates p<0.05, ^∗∗^ p<0.001, ^∗∗∗^ p <0.0001, ^∗∗∗∗^ p<0.00001, determined by two-tailed Student’ *t* test (D-K). MRT68921 abbreviated as MRT.

**Figure S3. Cells resistant to autophagy inhibition undergo anaplerosis and maintain basal respiration rates, related to Figure 2**

(A) Seahorse assay in FC1245 CTL and HCQ-R cells.

(B) Tracing map for [U-^13^C_5_] glutamine trace into TCA cycle.

(C-G) Abundance (left) and individual MIDs (right) for ⍺KG (C), Citrate (D), Fumarate (E), Malate (F), and Aspartate (G) in FC1199 CTL and HCQ-R cells following a 24h [U-^13^C_5_] glutamine trace.

Data are represented as a mean ± SEM. Panel A represents n=3 technical replicates representative of n=3 biological replicates. Panels C-G represents n=3 technical replicates. ^∗^ indicates p<0.05, ^∗∗^ p<0.001, ^∗∗∗^ p <0.0001, ^∗∗∗∗^ p<0.00001, determined by two-tailed Student’ *t* test (A and C-G) and two-way ANOVA with Sidak’s multiple comparison test. MRT68921 abbreviated as MRT.

**Figure S4. Resistance to autophagy inhibition causes a decrease in nucleotide pools and pyrimidine metabolism reprogramming, related to Figure 4**.

(A-B) Targeted metabolomics for nucleotide pools in FC1199 CTL and HCQ-R cells (A) or FC1199 CTL and MRT-R cells (B).

(C) *Dck, Tk1, Slc29a1, Slc29a2* and *Uck1* normalized mRNA expression in FC1199 CTL, HCQ-R and MRT-R cells.

(D-F) MTT cell viability assay in FC1199 CTL and HCQ-R cells treated with BQN (D) and BAY-2402234 (E), in FC1245 CTL and HCQ-R cells treated with BAY-2402234 (F) and in FC1199 CTL and MRT-R cells treated with BAY-2402234 (G) for 72h.

Data are represented as a mean ± SEM. Panels A-C represent n=3 biological replicates. Panel D-G represents n=3 technical replicates representative of n=3 biological replicates. ^∗^ indicates p<0.05, ^∗∗^ p<0.001, ^∗∗∗^ p <0.0001, ^∗∗∗∗^ p<0.00001, determined by two-tailed Student’ *t* test (A-B) and one-way ANOVA with Tukey’s post hoc test (C). MRT68921 abbreviated as MRT.

**Figure S5. Resistance to autophagy inhibiton increases sensitivity to pyrimidine analogs across different PDA cell lines**

(A-D) MTT cell viability assay and corresponding IC50 values in FC1245 CTL and HCQ-R cells treated with gemcitabine (A-B) and FTD-TPI (C-D) for 72h.

(E-F) MTT cell viability assay (E) and corresponding IC50 values (F) in Panc-1 CTL and HCQ-R cells treated with gemcitabine for 72h.

(G) Incucyte live cell imaging in FC1245 CTL and HCQ-R cells treated with gemcitabine (5nM) for 3 days.

(H-K) MTT cell viability assay and corresponding IC50 in FC1199 (H-I) anf FC1245 (J-K) CTL and HCQ-R cells treated with Triapine for 72h.

(L) Clonogenics assay in FC1199 CTL and HCQ-R cells treated with gemcitabine (5nM) or vehicle control with or without uridine (1mM). Images are representative of n= 3 biological experiments.

(M) Incucyte live cell imaging in FC1245 CTL and HCQ-R cells treated with gemcitabine (5nM) with or without deoxycytidine (100μM) for 3 days.

(N) Incucyte live cell imaging in FC1199 CTL and MRT-R cells treated with FTD/TPI (37.5nM) with or without uridine (1mM) for 3 days.

(O) Western blot for dCK protein and Tubulin in FC1199 CTL and HCQ-R cells KO for dCK with guide 1 (dCK1) or guide 2 (dCK2) and non-targeting (NT) control.

(P-U) MTT cell viability assay and corresponding IC50 values in FC1199 CTL, HCQ-R and HCQ-R drug-holiday cells treated with HCQ (P-Q), MRT68921 (R-S) and gemcitabine (T-U) for 72h.

(V-W) Images (V) and weights (W) of orthotopic FC1199 CTL and HCQ-R tumors from mice treated with gemcitabine or saline control (n=9-10/group).

Data are represented as a mean ± SEM. Panels A, C, E, G-H, J, M-N, P, R, and T represent n=3 technical replicates representative of n=3 biological replicates. Panels B, D, F, I, K, Q, S and U represent n=3 biological replicates. ∗ indicates p<0.05, ∗∗ p<0.001, ∗∗∗ p <0.0001, ∗∗∗∗ p<0.00001, determined by two-tailed Student’ t test (B, D, F, I, K, Q, S and U), two-way repeated measures ANOVA with Tukey’s post hoc test (G and M-N) and Kruskal-Wallis test (W). MRT68921 abbreviated as MRT.

**Figure S6: Autophagy inhibitors and pyrimidine analogues have a combinatory effect across different PDA cell lines, related to Figure 6**

(A) Correlation between ATG13 (left) or ULK1 (right) gene expression and gemcitabine sensitivity in PDA cells determined by single linear regression using Depmap data.

(B-D) Incucyte live cell imaging in FC1199 cells, (B) FC1245 cells or (C) Panc-1 cells treated with HCQ and/or gemcitabine as indicated for 3 days.

(E-F) Clonogenic assay in FC1245 cells treated with the combination of HCQ and gemcitabine or HCQ and FTD/TPI as indicated for 3 days.

(G-H) Incucyte live cell imaging in FC1245 cells treated with the combination of MRT68921 and gemcitabine or MRT68921 and FTD/TPI as indicated for 3 days.

(I) Clonogenic assay in FC1245 cells treated with the combination of MRT68921 and gemcitabine as indicated for 3 days.

(J-K) Incucyte live cell imaging in Panc-1 cells treated with the combination of MRT68921 and gemcitabine or HCQ and FTD/TPI as indicated for 3 days.

Data are represented as a mean ± SEM. Data on panels B-D, G-H and G-K represent n=3 technical replicates representative of n=3 biological replicates. Data on panels E-F and I are representative of n=3 biological replicates. ^∗^ indicates p<0.05, ^∗∗^ p<0.001, ^∗∗∗^ p <0.0001, ^∗∗∗∗^ p<0.00001, determined by two-way repeated measures ANOVA with Tukey’s post hoc test (B-D, G-H and G-K). MRT68921 abbreviated as MRT.

## EXPERIMENTAL MODELS AND SUBJECT DETAILS

### Cell Lines

All cell lines were maintained at 37°C and 5 % CO_2_. FC1199 and FC1245 cells were provided by the Tuveson lab and derived from the PDA tumor of a KPC male and female mouse, respectively. These cells were maintained in Dulbecco’s Modified Eagle Medium (DMEM; Thermo Fisher Scientific) with 10 % fetal bovine serum (FBS; Omega Scientific Inc.) and 1% Penicillin-Streptomycin (Pen/Strep; Thermo Fisher Scientific). Panc-1 cells were provided from the Engle lab and kept in Roswell Park Memorial Institute medium (RPMI 1640; Thermo Fisher Scientific) with 10 % FBS (Omega Scientific Inc.) and 1% Pen/Strep (Thermo Fisher Scientific). All cell lines were periodically monitored for mycoplasma contamination with the MycoAlert Mycoplasma Detection Kit (Lonza). FC1199, FC145 and Panc-1 cells resistant to HCQ were generated. The IC50 for HCQ (Sigma-Aldrich) was determined for each cell line using an MTT cell viability assay. Each cell line was treated with the IC50 for several weeks until little cell death was observed. A new IC50 was then determined, and cells were cultured with this new drug dose for several weeks. This process was repeated until cells reached a 5-10-fold resistance compared to CTL drug naïve cells kept in culture in parallel. Once this level of resistance was reached, the cells were kept in culture with drug in the media. FC1199 resistant to MRT68921 (MedChemExpress) were also generated using the same method.

### Animal Studies

10–12-week-old C57BL/6J male mice were purchased from The Jackson Laboratory. Mice were housed in filter-topped cages at room temperature (21°C) in the same room with a 12-hour light: dark cycle. They were fed a standard diet and had access to water ad libitum. 10–12-week-old male mice and 10-week-old female mice were used for FC1199 and FC1245 tumor implantation experiments, respectively. Mice were randomized onto treatment arms and all experimental protocols were approved and performed according to the Institutional Animal Care and Use Committee (IACUC) of the Salk Institute for Biological Studies.

## METHOD DETAILS

### MTT cell viability assay

PDA cells were plated in 96 well plates (1500-3000 cells per well) or 384 clear well plates (250-500 cells per well). The next day, cells were treated with different doses of the drug of interest in triplicate either manually or with the D300e digital dispenser (Tecan) 72h after, cell viability was measured using MTT assays (R&D Systems) according to the manufacturer’s protocol. The Infinite 200 pro M Plex microplate reader (Tecan) was used to measure the absorbance of all wells (including the blanks containing medium only) at 565nM for 180 seconds following a 2min shaking period. The IC50 values were calculated using Graph Pad Prism 10.1.1 (Graph Pad Software Inc).

### Incucyte live cell imaging

Live cell imaging was performed with the Incucyte S3 (Sartorius) at 10X magnification. FC1199, FC1245 (1500 cells per well) and Panc-1 CTL cells (5000 cells per well) as well as FC1199, FC1245 (2500 cells per well), Panc-1 HCQ-R cells (10 000 cells per well) and FC1199 MRT-R cells (2500 cells per well) were seeded in 96 well plates. These different cell numbers were used to account for differences in cell growth. Media was removed from all wells the next day and media containing the drugs of interest was added. The plates were then placed in the incucyte for 72h, with pictures taken every 4 hours. Cell confluency was determined using the Incucyte S3 software (Sartorius). The CellEvent Caspase 3/7 green reagent (Thermo Fisher Scientific) was used at a 1:1000 concentration to measure caspase 3/7 activity. The reagent was added at the same time as the drugs and green events were counted (optimized for each cell type) and reported relative to cell confluency.

### Clonogenic assay

500 cells were plated into 6 well plates. The next day, the media was removed and replaced with media containing drugs of interest or vehicle control. After 72 hours, the media was replaced with drug-free full media for another 5 days. Cells were then washed fixed and stained with crystal violet (Sigma-Aldrich).

### Measurement of Autophagic Flux by Ratiometric Flow Cytometry

FC1199 CTL, HCQ-R and MRT-R cells were transfected with the mCherry-EGFP-LC3 tandem, kindly gifted by Jayanta Debnath, and used for flow cytometric analysis. Cells were either (1) kept in normal media, (2) changed to EBSS starvation media (Sigma-Aldrich) for 24 hours, or (3) treated with the lysosomal inhibitor bafilomycin A1 (20 nM; Sigma-Aldrich) for 24 hours. Flow cytometry was performed with the BD FACSAria Fusion flow cytometer using 488 and 561nM lasers for green and red fluorophore excitation, respectively. The appropriate side/forward scatter profile was used to exclude non-viable cells. Autophagic delivery of the tandem constructs to the lysosome quenches the GFP signal, therefore cells undergoing autophagy are defined based on their mCherry/GFP fluorescence ratio. The gate to define cells undergoing autophagy was set based on cells treated with bafilomycin A1, a condition that represents cells with little or no autophagic flux. The bottom of the gate for each set of flow cytometry experiments was therefore set at the rightward base of the bafilomycin A1-treated curve such that 5% of bafilomycin A1-treated cells were included in the gate.

### RNA isolation and RNA sequencing

Total RNA was extracted from cells using the RNeasy Plus mini kit (Qiagen) according to the manufacturer’s recommendations. RNA concentration was assessed by Qubit Flex Fluorometer (Thermo Fisher Scientific, CA, USA). RNA integrity number was verified with Agilent 2200 TapeStation system (Agilent Technologies, Amsterdam, Netherlands). RNA-seq libraries (cDNA libraries from polyA mRNA) and sequencing were performed at The Razavi Newman Integrative Genomics and Bioinformatics Core Facility of the Salk Institute (RRID:SCR_014842 and SCR_014846). Libraries were prepared by using Illumina’s TruSeq RNA library Preparation kit according to manufacturer’s instructions (Illumina, San Diego, CA, USA). Libraries were pooled and sequenced using either the Illumina NovaSex 6000 or NovaSeq X Plus (Illumina, San Diego, CA, USA) with 50-bp paired-end read chemistry. Raw basecalls were demultiplexed to FastQ files using BCL2FastQ v v2.20.0.422 (Illumina, San Diego, CA, USA) and quality assessed using FastQC v0.11.8 (Babraham Bioinformatics, Cambridge, UK).

### Metabolite extraction and GC-MS analysis

At the conclusion of the tracer experiment, media was aspirated from all wells. Cells were rinsed on ice with 0.9% saline solution and lysed with 500μl of ice-cold methanol. After 1 min, 100 μl of water containing norvaline (1 μg/ml) was added to each sample and vortexed for 5 min and 50uL was plated on a 96 well plate aside for protein concentration assessment. The 450uL remaining of the lysates were centrifuged at 21,130*g* for 10 min at 4°C. The supernatant was transferred in a microvial (Thermo Fisher Scientific) before being evaporated with a speed-vacuum at 4°C. Dried polar metabolites were processed for gas chromatography–mass spectrometry (GC-MS) as previously described by Cordes and Metallo^72^. Briefly, polar metabolites were derivatized using a Gerstel MultiPur-pose Sampler (MPS 2XL). Methoxime–tert-butyldimethylchlorosi-lane (tBDMS) derivatives were formed by addition of 15 μl of 2%(w/v) methoxylamine hydrochloride (MP Biomedicals, Solon,OH) in pyridine and incubated at 45°C for 60 min. Samples were then silylated by addition of 15 μl of N-tert-butyldimethylsily-N-methyltrifluoroacetamide (MTBSTFA) with 1% tBDMS (RegisTechnologies, Morton Grove, IL) and incubated at 45°C for 30min. Derivatized samples were analyzed using gas chromatography–mass spectrometry (GC–MS) on a DB-35MS column (30 m × 0.25 mm internal diameter × 0.25 μm, Agilent J&W Scientific) installed in an Agilent 7890A gas chromatograph coupled to an Agilent 5975C mass spectrometer. Samples were injected at an initial oven temperature of 100 °C, held for 1 min, ramped at 3.5 °C/min up to 255°C, then at 15 °C/min up to 320 °C, and held for 3 min. Electron impact ionization was used, and the instrument scanned over the *m/z* range of 100–650.Metabolite quantification and isotopomer distribution analysis were performed using in-house MATLAB scripts, including correction for natural isotope abundance.

### High-resolution LC-MS/MS of polar metabolites

Polar metabolites were extracted from confluent six-well plates after growing in DMEM + 10% FBS + 1% penicillin streptomycin for 48 hours. At the conclusion of the experiment, media was aspirated, and the cells were rinsed with saline solution. Cells were lysed with 500 μl of ice-cold methanol and 200ul of water spiked with norvaline (1 μg/ml) and scraped into Eppendorf tubes. The tubes were vortexed for 5 min and centrifuged at 21,230g at 4°C for 5 min. Polar phase was dried under air, and the pellet was resuspended in 50 μl of acetonitrile and loaded onto LC-MS/MS as previously described^73^. Briefly, samples were loaded on to a Q Exactive orbitrap mass spectrometer with a Vanquish Flex Binary UHPLC system (Thermo Fisher Scientific) that was used with an iHILIC-(P) Classic, 150 by 2.1 mm, 5-μm particle,200 Å (Hilicon) column at 45°C. Five microliters of sample were injected. Chromatography was performed using a gradient of 20 mM ammonium carbonate, adjusted to pH 9.4 with 0.1% ammoniumhydroxide (25%) solution (mobile phase A) and 100% acetonitrile (mobile phase B), both at a flow rate of 0.2 ml/min. The LC gradient ran linearly from 80 to 20% B from 2 to 17 min and then from 20 to80% B from 17 to 18 min and then held at 80% B from 18 to 25 min.

### Targeted metabolomics for nucleotide pools

Cell pellets were extracted with the addition of 500 µL of chilled 80% methanol and 10 µL each of the internal standards’ glutamic acid-d5 and adenosine triphosphate-13C10, followed by cycles of 2 min sonication in an ice bath and 30 sec vortexing to resuspend cell pellets. Samples were next placed on dry ice for 30 min prior to centrifugation (14,000 x g, 4 C, 10 min). The supernatant was transferred to a new tube and dried under nitrogen gas. Samples were reconstituted in 100 µL of 20% methanol, vortexed 30 sec, and transferred to HPLC vials prior to LC-MS analysis for nucleotides. The remaining cell pellet underwent total protein concentration analysis on a Synergy H1 Hybrid reader (BioTek Instruments, Inc., Winooski, VT, USA). Nucleotides were analyzed on an Ultimate 3000 UHPLC system (Thermo Fisher, Waltham, MA, USA) coupled to a TSQ Quantiva mass spectrometer (Thermo Fisher) with separation performed on a Waters 4.6 x 100 mm, 3.5 µm XBridge Amide HILIC column (Waters Corporation, Milford, MA, USA). LC solvents included mobile phase A consisting of 95:5 water:acetonitrile, 20 mM ammonium hydroxide, 20 mM ammonium acetate, and mobile phase B consisting of 100 % acetonitrile. A gradient was used with a flow rate of 0.4 mL/min. Injection volume was 10 µL, the column oven temperature was set to 25 C. MS analyses were performed using electrospray ionization in negative ionization mode. Spray voltage was −3.5kV, ion transfer tube temperature of 325 C, and vaporizer temperature of 275 C. Multiple reaction monitoring (MRM) was performed. Skyline software v. 24.1.0.199^74^ was used to measure peak areas. Peak areas were normalized using internal standards and protein concentration.

### Seahorse assay

10,000 cells were plated in 96-well plates (XFe96 plates, Agilent) in technical replicates of 6-8 and incubated at 37°C overnight. The next day, the medium was changed to XF Assay Medium (Agilent) containing 10 mM glucose, 1 mM pyruvate, and 2 mM glutamine and the cells were incubated at 37°C for one hour in a non-CO_2_ incubator. The Seahorse XF mitochondrial stress test (Agilent) was performed according to manufacturer’s instructions and oxygen consumption rates (OCR) were measured with a Seahorse XFe96 extracellular flux analyzer (Agilent). Briefly, basal OCR was measured prior to the addition of oligomycin (0.5μM), followed by 3 more OCR measurements and then carbonilcyanide p-triflouromethoxyphenylhydrazone (FCCP) (2μM) was added. Three more OCR measurements were taken before Rotenone/Antimycin-A (0.5μM) was added followed by three final OCR measurements. All data was analyzed in Wave software and normalized to confluency (assessed by a scan-on-demand in the Incucyte performed immediately after the mito stress test). The parameter values were calculated in Wave as follows: Basal respiration – Last rate measurement before the first injection subtract the non-mitochondrial respiration rate (minimum rate measurement after Rotenone/Antimycin-A injection), spare respiratory capacity – maximal respiration (maximum rate measurement after FCCCP injection subtract the non-mitochondrial respiration rate) subtract basal respiration.

### Western Blotting

Whole cell lysate samples were washed with cold PBS and harvested on ice with RIPA buffer and 1X of protease inhibitor cocktail (Roche) added just before use. Lysates were sonicated before being spined down at 20 000xg for 15min at 4C. The pellet was discarded, and supernatant was kept. Protein concentration was determined with a Bradford assay and was measured relative to a bovine serum albumin (BSA) standard curve. Western blotting was performed using standard methods including protein separation on SDS-PAGE 1.5mm mini gels in running buffer at 100V for 2hrs, followed by transfer to PVDF membranes in transfer buffer using a semi-dry transfer apparatus at 15V for 70 minutes. Membranes were blocked in 5% milk for 1hr at room temperature with gentle rocking, washed twice in 1X TBST, and then incubated overnight at 4°C with gentle rocking in primary antibodies. Membranes were then washed three times in TBST and incubated for 1hr at room temperature with gently rocking in secondary antibodies followed by 3 more TBST washes. Membranes were developed with Immobilon Western chemiluminescent HRP substrate (Sigma-Aldrich) and analyzed on the ChemiDoc imaging system (Bio-Rad). Antibodies are listed in the Key Resources Table.

### Immunofluorescence on cell cultures

Cells were grown on glass cover slips (Neuvitro) disinfected with 30min UV irradiation before being treated with Poly-L-lysine (Cell Biologics) for 3 hours. The cells were fixed with 10% formalin at room temperature for 15min and washed three times with PBS (Thermo Fisher Scientific). Cells were then permeabilized with 0.1% Triton X-100 (Research Products Internation Corp.) for 5min at room temperature, followed by another three PBS (Thermo Fisher Scientific) washes. After a 30 min block in a mixture of 5% goat serum (Vector Laboratories), 0.3 M glycine (Thermo Fisher Scientific) and PBS (Thermo Fisher Scientific) at room temperature, anti-TOMM20 (Abcam) and anti-PDH (Abcam) primary antibodies diluted in blocking buffer were added to incubate overnight at 4°C. Following three PBS washes, cells were incubated with Alexa Fluor conjugated secondary antibodies (Thermo Fisher Scientific) at room temperature for 45min protected from light. After three other PBS washes, the cover slips were mounted onto glass slides with mounting medium (Vector Laboratories) and left to dry overnight protected from light. The slides were then imaged with a confocal laser-scanning Olympus FV3000 with a 60X objective. Z-stacked images were taken and then displayed as max-projections.

### Electroporation with RNPs

The CRISPR/Cas9 system was used to generate DHODH KO and non-targeting (NT) control FC1199 cells. Electroporation was performed using the 4D-Nucleofector system (Lonza Bioscience) with the P3 Primary Cell 4D-Nucleofector X Kit S (Lonza Bioscience), following the manufacturer’s instructions. sgRNA were obtained from Synthego and reconstituted in TE buffer at 0.5nM/μL. To form RNPs for each condition, 0.5nM of sgRNA was incubated at room temperature for 10min with 6μg of Cas9 (PNA Bio), previously resuspended in a 20% glycerol buffer at 5μg/ml. Two million FC1199 cells were mixed with the RNP complexes and transferred to a well of a Nucleocuvette Strip. The strip was then electroporated on the Lonza 4D electroporation system with pulse code CZ167 (Cell Type Program MEF, mouse). Knock-out was confirmed by western blot one week post electroporation.

### Pharmacotyping in hPDOs and response correlation with gene expression

We used previously published methods and organoids^58^. Briefly, PDAC organoids were plated in 384-well plates at the density of 1,000 cells per well in 20μL media. Drugs were dispensed in each well using the D300e Digital Dispenser (Tecan). 5-days after treatment, cell viability was measured using a luminescent ATP-based assay (CellTiter-Glo, Promega) and Synergy H1 plate reader. To assess hPDO response correlation with gene expression, we performed a Pearson correlation of gemcitabine sensitivity (area under the curve from Tiriac et al.^58^, low AUC indicates sensitivity to gemcitabine) and the Autophagy gene signature.

### Mouse treatments

2000 FC1199 CTL and 2000 or 10 000 FC1199 HCQ-R cells were injected orthotopically in the pancreas of 10-week-old C57BL/6J male mice (Jackson) in 30uL of a 1:1 serum-free DMEM media (Thermo Fisher Scientific) and Matrigel (Corning) mixture. Eight days after the injection, mice received gemcitabine (5mg/kg, IP, every 3 days; McKesson Medical-Surgical Inc.) or saline control (Pfizer). Mice were euthanized 21 days after cancer cell injections. 2000 FC1245 CTL and HCQ-R cells were injected orthotopically in the pancreas of 10-week-old C57BL/6J female mice as described above. Gemcitabine treatment (5mg/kg, IP, every 3 days; McKesson Medical-Surgical Inc.) or saline (Pfizer) control was initiated 12 days later. Mice were euthanized 22 days after cancer cell injections. 1000 drug-naïve FC1199 cells were also injected orthotopically in the pancreas of 12-week-old C57BL/6J male mice as described in this section. The mice received SBP-1750 (30mg/kg, PO, daily) or vehicle control, and gemcitabine (10mg/kg, IP, every 3 days; McKesson Medical-Surgical Inc.) or saline (Pfizer) control. Mice were euthanized 28 days after the orthotopic surgery, and 2-3 hours after the last drug treatment.

### Histology

Mice were euthanized and tissues were fixed in 10% neutral buffered formalin (Epredia). All sample preparation and staining procedures were then performed at HistoWiz Inc., using HistoWiz’s Standard Operating Procedure, and a fully automated workflow. Samples were processed, embedded in paraffin, and sectioned at 4 μm. Slides were stained by a standard H&E staining protocol (Histowiz, Long Island City NY). The slides were then dried, and film-coverslipped using a TissueTek-Prisma and Coverslipper (Sakura). Whole-slide scanning (40×) was performed on an Aperio AT2 (Leica Biosystems).

### Quantification and Statistical Analysis

Statistical analyses and graphs were performed using Graph Pad Prism 10.1.1 (Graph Pad Software Inc). A two-tailed Students t-test was used for comparisons between two groups. For analyses involving more than two groups, either a one-way ANOVA followed by Tukey’s post hoc test (for a single variable) or a two-way ANOVA with Tukey’s post hoc test (for multiple variables) was performed. For non-normally distributed data, a Mann-Whitney test or a Kruskal-Wallis test were performed for comparisons between two groups and more than two groups with a single variable, respectively. Graphs were made with Graph Pad Prism 10.1.1 (Graph Pad Software Inc). with the error bars representing mean with standard deviation. All group numbers and explanation of significant values are presented within the figure legends. ^∗^p < 0.05; ^∗∗^p < 0.01; ^∗∗∗^p < 0.001; ^∗∗∗∗^p < 0.0001.

