## Supplementary material for "Leveraging autophagy and pyrimidine metabolism to target pancreatic cancer": Key Resource Table

**Key resources table**

| REAGENT or RESOURCE | SOURCE | IDENTIFIER |
|---|---|---|
| Antibodies | | |
| Goat Anti-Mouse IgG Antibody, Alexa Fluor 488 Conjugated | Thermo Fisher Scientific | Cat# A11005; RRID:AB_2534073 |
| Goat Anti-Rabbit IgG Antibody, Alexa Fluor 488 Conjugated | Thermo Fisher Scientific | Cat#A11008; RRID:AB_143165 |
| Anti-Mouse IgG | Cell Signaling Technology | Cat#53484; RRID:AB_2799435 |
| Anti-Rabbit IgG | Cell Signaling Technology | Cat#7074; RRID:AB_2099233 |
| ATG13 | Cell Signaling Technology | Cat#13273; RRID:AB_2798169 |
| β-ACTIN | Sigma-Aldrich | Cat#A5441; RRID:AB_476744 |
| CAD | Cell Signaling Technology | Cat#11933; RRID:AB_2797772 |
| DCK | Abcam Biochemicals | Cat#ab186128; RRID:AB_3696024 |
| DHODH | Proteintech | Cat#14877-1-AP; RRID:AB_2091723 |
| GAPDH | Cell Signaling Technology | Cat#2118; RRID:AB_561053 |
| PCX | Proteintech | Cat#16588-1-AP; RRID:AB_1851513 |
| PDHA1 | Abcam | Cat#ab110333; RRID:AB_10862029 |
| SLC29A1 (ENT1) | Proteintech | Cat#11337-1-AP; RRID:AB_2190784 |
| TK1 | Proteintech | Cat#15691-1-AP; RRID:AB_2203613 |
| TOMM20 | Abcam | Cat#ab78547; RRID:AB_2043078 |
| TPK1 | Proteintech | Cat#10942-1-AP; RRID:AB_2205647 |
| α-TUBULIN | Cell Signaling Technology | Cat#2144; RRID:AB_2210548 |
| Chemicals, peptides, and recombinant proteins | | |
| [3-$^{13}C_1$] glucose tracer | Cambridge Isotopes | CLM-1393 |
| [U-$^{13}C_6$] glucose tracer | Cambridge Isotopes | CLM-1396 |
| [U-$^{13}C_6$] glutamine tracer | Cambridge Isotopes | CLM-1822 |
| 10% Neutral buffered formalin | Epredia | 5725 |
| 5-FU | Selleck Chemicals | S1209 |
| Bafilomycin A1 | Sigma-Aldrich | 196000 |
| Bovine Serum Albumin (BSA) | Sigma-Aldrich | A9647 |
| Chloroform | Sigma-Aldrich | 366927 |
| Crystal Violet | Sigma-Aldrich | C6158 |
| Cytarabine | Thermo Scientific Chemicals | 449560010 |
| Deoxycytidine | Thermo Scientific Chemicals | J63062.06 |

| | | |
|---|---|---|
| Dialized Fetal Bovine Serum | Thermo Fisher Scientific | 26400044 |
| DMEM | Thermo Fisher Scientific | 10013CV |
| DMEM – glucose free | Thermo Fisher Scientific | 11966025 |
| DMEM – glutamine free | Thermo Fisher Scientific | 10313021 |
| DMSO | Corning | 25950CQC |
| EBSS | Sigma-Aldrich | E2888 |
| Fetal Bovine Serum | Omega Scientific | FB-01 |
| Hydroxychloroquine | Sigma-Aldrich | H0915 |
| Gemcitabine HCl | R&D Systems | 3259 |
| Gemcitabine HCl (Pharma grade) | McKesson Medical-Surgical | 1125820 |
| Glycine | Thermo Fisher Scientific | BP3811 |
| Goat serum blocking solution | Vector Laboratories | S-1000 |
| L-Glutamine (α-$^{15}$N) | Cambridge Isotopes | NLM-1016 |
| Matrigel | Corning | 356231 |
| Methanol | Sigma-Aldrich | 34860 |
| MRT68921 | MedChemExpress | HY-100006A |
| Opti-MEM | Thermo Fisher Scientific | 11058021 |
| Oxaliplatin | Adipogen Life Sciences | AG-CR1-3592 |
| Paclitaxel | Cayman Chemical | 10461 |
| PageRuler Prestained Protein Ladder | Thermo Fisher Scientific | 26616 |
| PBS | Thermo Fisher Scientific | 141190144 |
| Penicillin-streptomycin | Thermo Fisher Scientific | 15140122 |
| RPMI | Thermo Fisher Scientific | 10040CV |
| Saline solution | Pfizer | 04094888 |
| SBP-1750 | Sanford Burnham Prebys | n/a |
| SN38 | Selleck Chemicals | S4908 |
| Thymidine | Sigma-Aldrich | T9250 |
| Triapine | Selleck Chemical | S7470 |
| Triton X-100 | Research Products International Corp | 1110361L |
| Trifluridine-tipiracil (TAS-102; FTD/TPI) | MedChemExpress | HY-16478 |
| Trypsin-EDTA (0.25%), phenol red | Thermo Fisher Scientific | 25200114 |
| Tween 20 | Sigma-Aldrich | P9416 |
| Tween 80 | Thermo Fisher Scientific | 28329 |
| Ultra-pure water | Invitrogen | 10977015 |
| Uridine | Sigma-Aldrich | U3003 |
| XF Assay DMEM Medium | Agilent | 103680 |

| | | |
|---|---|---|
| Critical commercial assays | | |
| Seahorse XF Mitochondrial Stress Test | Agilent | 103015–100 |
| MTT Cell Proliferation/Viability Assay | R&D Systems | 4890-050-K |
| MycoAlert Mycoplasma | Lonza | LT07-318 |
| P3 Primary Cell 4D-Nucleofector™ | Lonza | V4XP-3032 |
| Protease Inhibitor Cocktail | Roche | 11873580001 |
| RIPA | Sigma-Aldrich | 20-188 |
| RNeasy Plus Mini Kit | Qiagen | 74134 |
| DC Protein assay | Bio-Rad | 5000116 |
| Pierce BCA Kit | Thermo Fisher Scientific | 23227 |
| P3 Primary Cell 4D-Nucleofector X Kit S | Lonza | V4XP-3032 |
| Immobilon Western Chemiluminescent HRP Substrate | Sigma-Aldrich | WBKLS0500 |
| Deposited data | | |
| RNA sequencing | | GEO submission pending (currently provided as supplemental table 1) |
| Experimental models: Cell lines | | |
| Mouse: FC1199 | Laboratory of David Tuveson, CSHL | n/a |
| Mouse: FC1245 | Laboratory of David Tuveson, CSHL | n/a |
| Human: MIA PaCa-2 | Laboratory of Dannielle D. Engle, Salk Institute | n/a |
| Experimental models: Organisms/strains | | |
| Mouse: C57BL/6 | Jackson | Cat#000664; RRID:IMSR_JAX:000664 |
| Recombinant DNA | | |
| pBABE-puro-mCherry-EGFP-LC3B | Jayanta Debnath | Addgene plasmid # 22418[83] |
| Software and algorithms | | |
| Biorender | https://www.biorender.com/ | |
| FlowJo 10.10.0 | Becton, Dickinson and Company (BD) | |
| GraphPad Prism 10.2.1 | GraphPad | |
| ImageJ | Scheinder et al. 2012[84] | |
| MATLAB R2022a | MathWorks | |
| QuPath 0.4.3 | QuPath | |
| Other | | |
| CellEvent Caspase3/7 Green Detection Reagent | Thermo Fisher Scientific | C10423 |
| Cas9 recombinant protein | PNA Bio | CP02 |
| Guide RNA – *Atg7* (mouse) | Synthego | GCAGCAGUGGGCAGGCGUGG |

| | | |
|---|---|---|
| Guide RNA – *Dck* (mouse) guide #1 | Synthego | UCUCUUGUACAACAGCUGCU |
| Guide RNA – *Dck* (mouse) guide #2 | Synthego | UAUCCUAAAGCAAGCCUCUG |
| Guide RNA – *Dhodh* (mouse) | Synthego | GAUGCAGCCAUCAUCCUUGG |
| Microvial | Thermo Fisher Scientific | 6ESV9-04PP |
| XFe96 plates | Agilent | 103794 |
| Glass cover slips | Neuvitro | GG121.5OZ |
